# Comprehensive analysis of structural and sequencing data reveals almost unconstrained chain pairing in TCRαβ complex

**DOI:** 10.1101/693630

**Authors:** Dmitrii S Shcherbinin, Vlad A Belousov, Mikhail Shugay

## Abstract

Antigen recognition by T-cells is guided by the T-cell receptor (TCR) heterodimer formed by α and β chains. A huge diversity of TCR sequences should be maintained by the immune system in order to be able to mount an effective response towards foreign pathogens, so, due to cooperative binding of α and β chains to the pathogen, any constraints on chain pairing can have a profound effect on immune repertoire structure, diversity and antigen specificity. By integrating available structural data and paired chain sequencing results we were able to show that there are almost no constraints on pairing in TCRαβ complexes, allowing naive T-cell repertoire to reach the highest possible diversity. Additional analysis reveals that the specific choice of contacting amino acids can still have a profound effect on complex conformation. Moreover, antigen-driven selection can distort the uniform landscape of chain pairing, while small, yet significant, differences in the pairing can be attributed to various specialized T-cell subsets such as MAIT and iNKT T-cells, as well as other putative invariant TCRs.

## Introduction

The process of somatic recombination can produce an immense diverse repertoire of TCR α and β chain sequences in human, having a theoretical bound on the number of unique variants of 10^10^ for a single chain [1]. The effective T-cell diversity is thus only limited by the total number of T-cells in human that is ∼10^11^ [1] and potential pairing preferences between α and β chains. The size of foreign peptide pool to be recognized by T-cells can reach ∼10^12^ variants for all possible 9-mers presented by HLA class I. So, as both TCR α and β chain is required to recognize an antigen [2] and a certain degree of cross-reactivity is needed to be able to form an efficient immune response [3], one would expect the immune system to aim at producing the highest possible number of α and β chain combinations in order to ensure optimal recognition of newly encountered pathogens.

Early estimates of the extent of pairing in TCRαβ complex [4] indicate that around 25 distinct α chains are paired to the same β chain in human T-cell repertoire. However, given the limited amount of unique TCRαβ clones that can be obtained via combinatorial single-chain high-throughput sequencing methods (∼10^4-5^ in PairSEQ datasets [5] or by frequency-based pairing [6]) and even smaller typical yield of single-cell methods (∼10^3-4^ cells according to 10x Genomics^®^ dataset compendium [7]), it is nearly impossible to directly quantify and enumerate the range of possible αβ pairings as recapturing the same α or β chain sequence is highly unlikely event for naive T-cells. The latter suggests that the exploration of pairing preferences should be performed indirectly by using statistical models to extrapolate consistent patterns observed in available TCRαβ repertoire data. For example, a recent study [8] uses statistical modelling to show that there are certain subtle (yet significant) biases in αβ pairing at the nucleotide level that stem from the genome organization of α and β loci and intrinsic biases of the V(D)J rearrangement process. Certain biases of αβ pairing were also reported to be related to CD4/CD8 T-cell differentiation [9].

In the present study, we utilize structural data to infer the set of contacting residues between α and β chains that should convey all pairing information in the absence of antigen-driven selection. We deliberately exclude the middle portion of CDR3 sequences that are involved in antigen recognition and are therefore subject to a multitude of selection biases.

We report a conserved pattern of TCRαβ contacts between human and mouse, some of which involve highly conserved residues so that no information can be extracted from them. Frequentist inference of pairing preferences obtained from high-throughput sequencing data revealed certain residues that are paired significantly more often than expected by chance. Bayesian analysis, however, shows that these residues are rarely encountered and there is insufficient information for building an accurate αβ pairing predictor, leading to a conclusion that the pairing is almost random. Finally, deeper analysis of available datasets suggests that pairing preferences can be heavily altered by antigen-driven selection and some αβ chain interactions encode invariant T-cell subsets.

## Materials and methods

### TCR:pMHC structural data analysis

We manually queried Protein Data Bank [10] for TCR:peptide:MHC complex structures. In total, 170 structures for both human and mouse were downloaded. All these structures were further cleaned up using the PyMOL software (Schrödinger, LLC) as follows: only one copy of each of the chains forming the complex (TCRα, TCRβ, peptide, MHCα and MHCβ/β2microglobulin) was left and all auxiliary proteins in case of several models; ions and ligands were deleted. After cleanup all structures were spatially superimposed according to the TCRα-TCRβ chain pair pose.

In order to summarize contact frequency at TCR α and β chain residues, we have set up a global indexing of residues across all possible V-J rearrangements. IMGT numbering [12] was used for the V gene residue indexing, with Cys anchor residue of the CDR3 having an index of 104^th^ and preceding residues indexed according to IMGT alignment with gaps for a given V allele. We have included Cys104 and the following 3 residues (105-107) of V germline part of CDR3 and last 4 residues (108-111) of the J germline part, thus 111^th^ residue in our indexing system is the Phe/Trp anchor of J gene (part of the ‘[FW]GXG’ motif). We included 4 flanking CDR3 residues in the indexing based on our observation that they have almost fixed spatial position in CDR3 loop and rarely contact the antigen.

Contacting residues between TCR α and β chains were selected if they have C_α_ atom distance closer than 15Å. We have also required a direct contact with a distance less than 5Å between closest atoms to be present for a given residue index pair (according to IMGT-based indexing introduced in previous section) for it to be considered in the analysis.

### PairSEQ αβ repertoire sequencing data

Data from a paired TCRαβ receptor sequencing experiment (hereafter termed PairSEQ) was obtained from the previously published study of Howie *et al*. [5]. We have pooled datasets from ‘Experiment 1-5’ provided with the study. The dataset was converted to FASTA format preserving TCR pairing information and processed by the MIXCR software to produce V and J gene and CDR3 nucleotide and amino acid sequence calls.

For some of our analysis focused on VαJαVβJβ assignments, we have summarized the data according to V and J gene call scores. We weighted each of the potential V-J assignments based on their score, for example, if we detect a TRBV-X with MIXCR match score of 200.0 and TRBV-Y with a score of 100.0 coupled with TRBJ-Z represented by 3 reads, we produce two distinct TCR clonotypes: TRBV-X:TRBJ-Z with a frequency of 2.0 reads and TRBV-Y:TRBJ-Z with a frequency of 1.0 read.

### Bayes network analysis

We used BNLearn R package [11] to infer Bayes Networks of amino acid probability distributions at TCR α and β chain residues. A hill-climbing (*hc*) and a hybrid algorithm (*rsmax2*) was used for the network construction with AIC score as an objective. After fitting to the data, inferred Bayes Networks were used for scoring TCRαβ complexes based on computed Log Likelihood (LL) values for instances. We have also heavily relied on the ability to both whitelist and blacklist an edge in order to decouple intra- and inter-chain contact networks.

### Specific TCR data analysis

Data on T-cell receptor sequences with known antigen specificity was downloaded from the VDJdb database (vdjdb.cdr3.net) and filtered to include only records where both α and β chain sequences are known. Only antigens with at least 30 paired-chain records were selected for further analysis.

### Source code and availability

An R markdown notebook with code for the analysis described in the manuscript together with corresponding datasets is available via GitHub at https://github.com/antigenomics/trab-pairing-study.

## Results

### Contact maps of TCRαβ complexes

We started our analysis with exploring contact frequencies of α and β chain residues of known TCR:peptide:MHC complexes available via the PDB database for both human and mouse. Next, we proofread and corrected the complexes and mapped Variable (V) and Joining (J) gene sequences to produce complementarity determining region (CDR)/framework region (FR) markup and enumerate the residues of TCR chains (see **Materials and Methods** section).

We utilized IMGT numbering [12] to index residues and bring all sequences to a uniform coordinate set. Additionally, we discarded the middle part of the CDR3 region as it is mostly involved in TCR:antigen recognition and forms an extremely flexible omega loop that doesn’t allow meaningful indexing across distinct TCRs.

The contact residues were defined based on C_α_ atom distance in order to compensate for the presence of distinct side chains that may not be fully covered by the available structural data (see **Materials and Methods** section). Resulting contact maps (**Figure 1**) show conservation between human and mouse and were highly symmetric: they feature contacts between FR1 and FR2 regions, FR2:FR2 contacts, FR2:CDR3 flank contacts and contacts between CDR3 flanks of different chains. There also is a visible non-symmetric contact region between the start of FR3 of β chain and J part of CDR3 of α chain.

**Figure 1.**
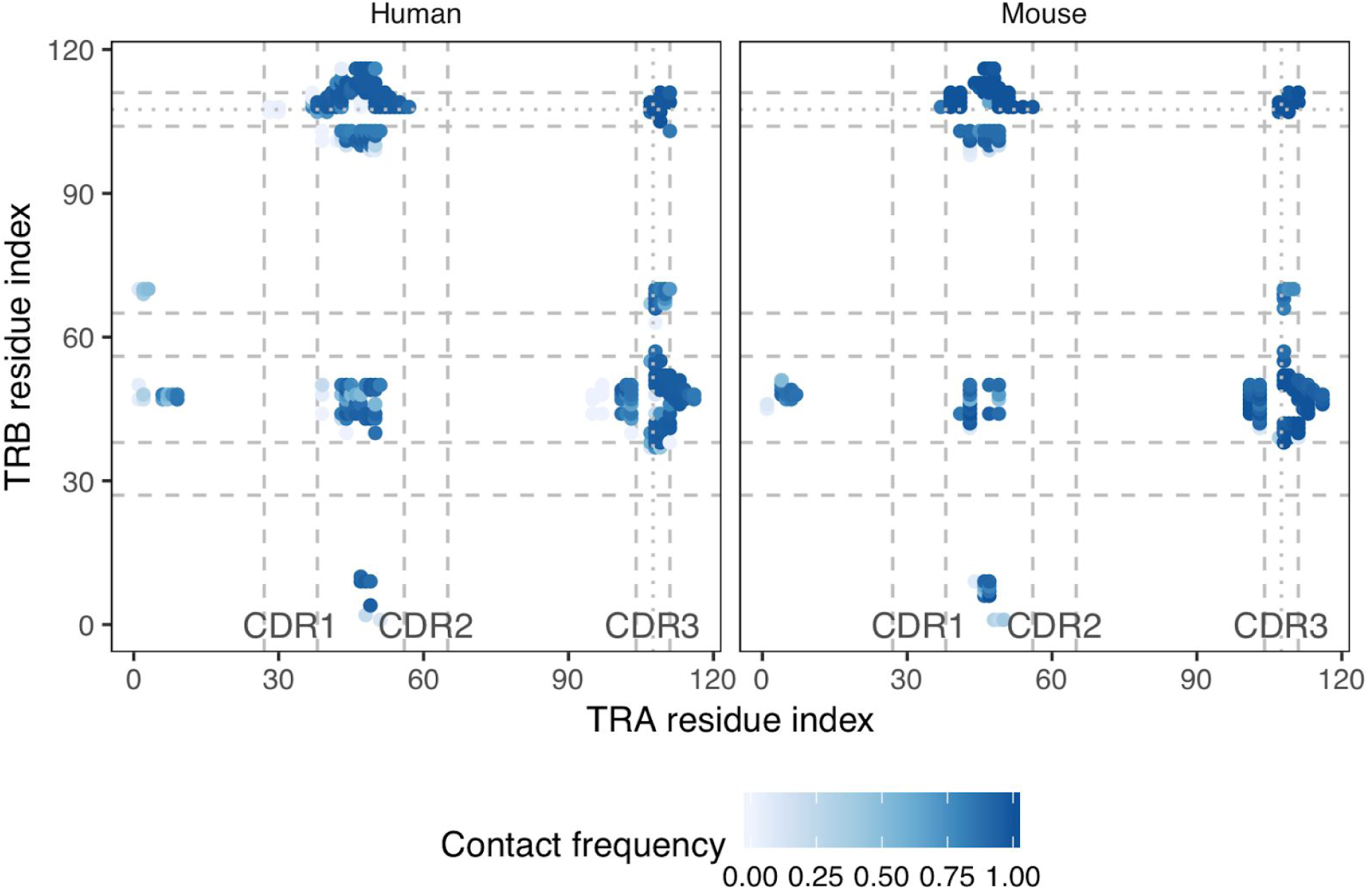
Contact map of TCRαβ complexes. A heatmap of inter-chain residue contact frequencies observed in *n = 131* human and *n = 39* mouse TCR:peptide:MHC complexes. Contacts were counted based on a C_α_ distance threshold of 15Å, for a residue pair to be included in the heatmap, it should directly contact a pair of atoms closer than 5Å in at least one complex. CDR regions are shown with dashed lines, excluded middle portion of CDR3 is shown with dotted line.

### Sequence conservation at contacting residues

We further explored the conservation of various residues of α and β chains and compared them to the contacts frequency that involved a given residue and any residue from the other chain (**Figure 2**). Contact residues can occur both in highly conserved and in less informative regions of V and J genes, and weighted conservation scores that account for differential V and J usage in observed repertoires didn’t change the picture (**Figure 2a**).

**Figure 2.**
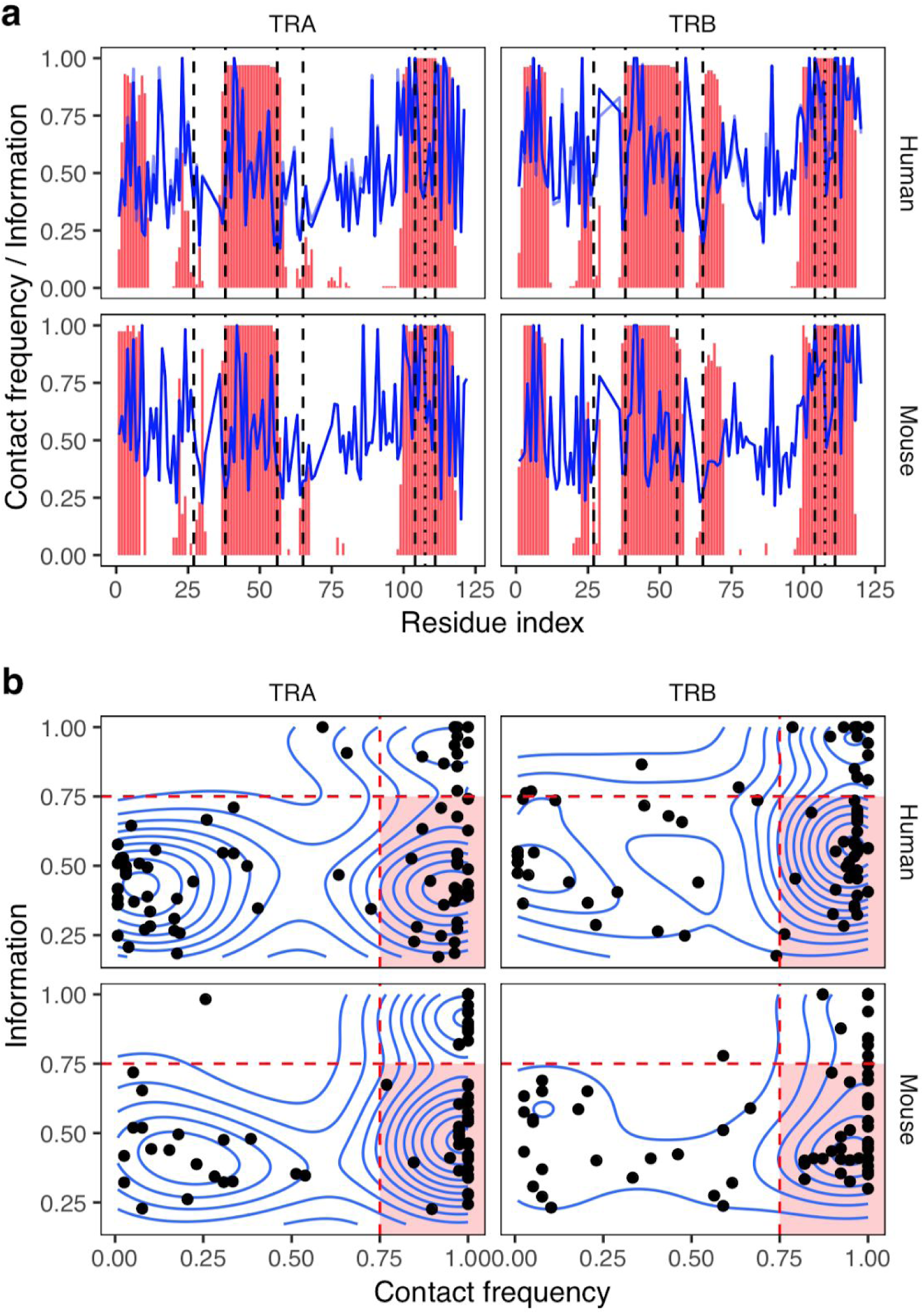
Information content and contact frequency of TCR chain residues. **a**. Contact frequency (red bars) and information content (blue line) across α and β chains. For human, transparent blue line shows information content computed from residue frequencies weighted by corresponding V and J gene usage in donor repertoires. Dashed lines show FR-CDR boundaries. b. Scatter plot represents the probability of being involved in inter-chain contact (x axis) and information content of amino acid distribution (y axis) for individual α and β chain residues. Region with a selection of frequently contacted residues (*x* > *0*.*75*) that are not too conservative (*y* < *0*.*75*) is highlighted with a red background.

While there is little correlation between residue conservation and contact frequency, scatter plot of these two variables for various residues show the presence of three groups of residues for both chains in human and mouse (**Figure 2b**). Almost all residues that are not frequently contacted are not well-conserved, on the other hand, there are two groups of residues involved in inter-chain contacts with high and low conservation. In our further analysis we focused only on the latter (highlighted area in **Figure 2b**), as they are both involved in direct inter-chain contacts and have enough variability in their amino acid content required to study chain pairing preferences.

### Statistics of amino acid pairing preferences at contact residues

In order to study the (non)randomness of chain pairing in TCRαβ complex we used a large dataset of TCR clones reported in the PairSEQ study [5] (see **Materials and Methods** section). We calculated the number of times a given amino acid pair is observed at a given contact (residue pair) for the set of residues selected as described in the previous section. We next normalized the resulting amino acid frequency matrices for each residue pair to calculate expected counts under the hypothesis of random amino acid pairing at a given contact.

The logarithm of the ratio of observed to expected counts of amino acid pairs is plotted in **Figure 3**. Ratios higher than 2.0 or less than 0.5 are only present for rare amino acid pairs with frequencies less than 10^3^ out of ∼10^5^ total observations, in line with the growing variance of the ratio for smaller counts. The latter suggests that there is little, if any, pairing preference and this pairing preference is limited to rare TCR clones.

**Figure 3.**
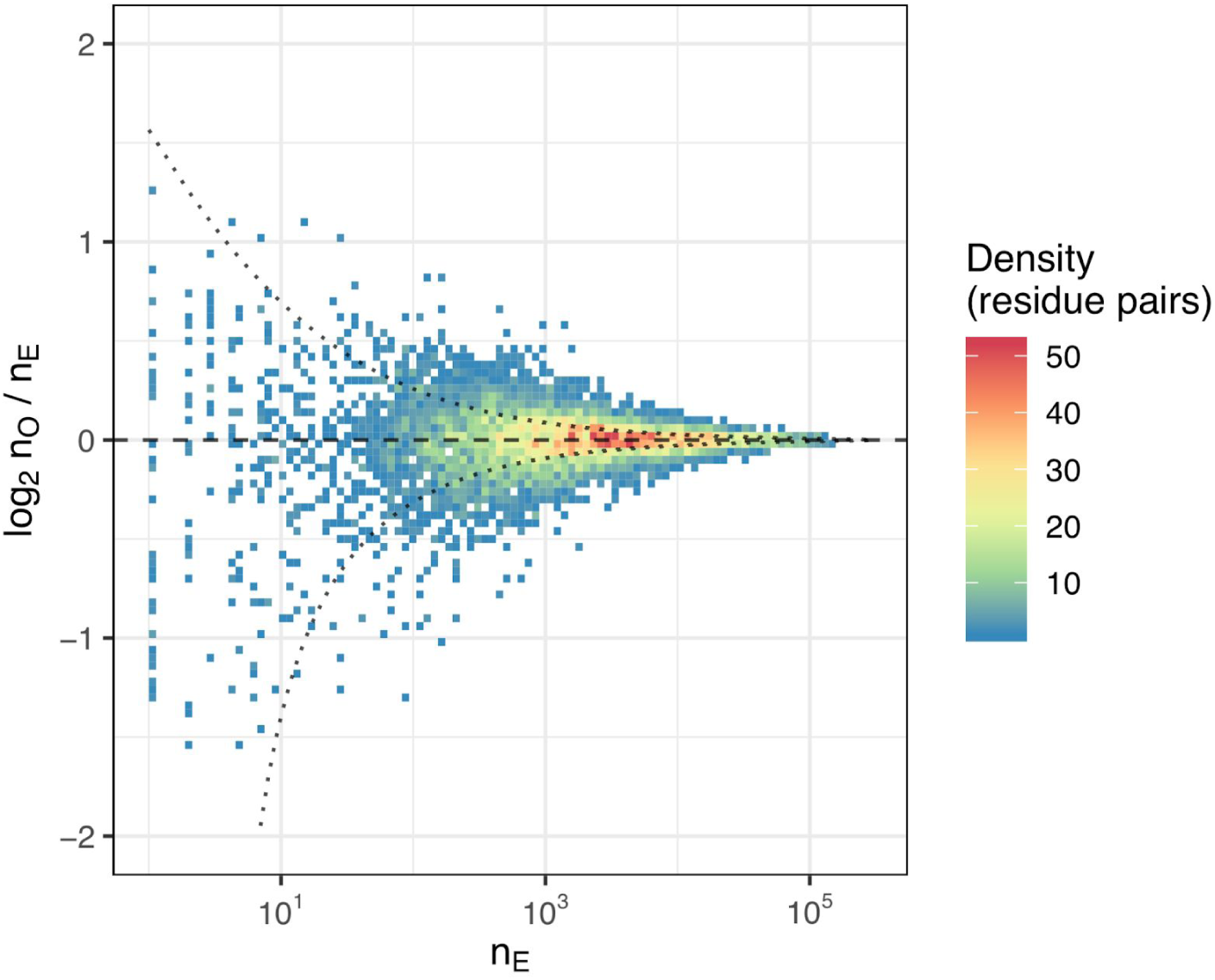
Amino acid statistics at contacting residues. Scatterplot showing n_E_, the number of times a given amino acid pair is expected to be present at a given inter-chain contact based on amino acid frequencies at separate chains, plotted versus the logarithm of the ratio of the observed amino acid pair count n_O_ to the expected count n_E_. Each point represents an inter-chain contact, i.e. at selected pairs of residue indices. The plot is based on n_T_=261,493 unique VαJαVβJβ combinations extracted from the PairSEQ dataset. Dotted lines show 95% confidence interval for the n_O_ / n_E_ ratio assuming Normal distribution with standard deviation computed from Binomial distribution approximated by n_E_ and n_T_.

We computed *χ*^2^ and mutual information (MI) values for amino acid frequency matrices, the relation between MI and both the conservation of contacting residues and contact frequency for a given residue pair. As can be seen in **Supplementary Figure S1a**, *χ*^2^ and MI values are in perfect agreement, yet one can note that the absolute value of MI < 0.003, meaning that while there are highly significant associations between contacting amino acids, the information carried by them is relatively small in agreement with observations from the previous paragraph. Highest MI scores are reached for the residue pairs with least conservation (**Supplementary Figure S1b**). There is little correlation between MI and residue pair contact frequency (**Supplementary Figure S1c**), suggesting the presence of indirect correlations.

An example of such indirect correlation between amino acid profiles is given in **Supplementary Figure S2a**, where the top 5 α chain residues that contact with β_101_ by their MI score were listed. Notably, residues α_47_ and α_43_ have similar MI scores yet the frequency of their contacts with β_101_ is substantially different, 95% and 11% respectively. As can be seen from amino acid frequency matrices in **Supplementary Figure S2b**, the amino acid preference profile is nearly the same for Tryptophan of α_47_ and Histidine of α_43_. On the other hand, there is only a single V gene that had both amino acids at given positions, as highlighted in **Supplementary Figure S2c**. Thus, one is able to assume that Tryptophan at α_47_ is directly contacted by β_101_ based on the contact frequency difference, while the high score of Histidine of α_43_ is merely an artifact arising from the linkage of α_47_ and α_43_ in a single V gene.

### Bayesian analysis reveals that pairing in TCRαβ complexes is almost random

In order to assess the actual information carried in inter-chain residue contacts that is relevant to TCRαβ chain pairing preferences we built a Bayes network (BN, see **Materials and Methods** section) of amino acid contact frequencies at residues that are frequently in contact between chains, as highlighted in **Supplementary Figure S2b**. BN also helped us to resolve indirect correlations shown in **Supplementary Figure S2c** and discussed in the previous section.

The BN construction for the entire PairSEQ dataset with no restriction of edges resulted in the structure shown in **Supplementary Figure S3**. In this case, a simple Hill-Climbing algorithm was used with a metric to optimize being Akaike Information Criterion (AIC, used in all BN learning throughout this section). Notably, only two edges connecting α and β components were observed: β_101_→α_43_ and α_46_→β_10_. The latter suggests that there is little information in inter-chain pairing, in line with the previous observations.

We therefore built a BN with only inter-chain contacts selected as described in the previous sections, blacklisting all intra-chain contacts as they contained several orders of magnitude more information that could disturb the assignment of inter-chain edges (**Supplementary Figure S4**). The positioning of inter-chain contacts that we learned is shown in **Supplementary Figure S5**. Notably, this contact map shows far fewer residues with multiple contacts, suggesting that BN helped to resolve ambiguous cases with indirect correlation as expected. An example of conditional probability matrix for α_101_ with both same chain parent α_55_ and an inter-chain interaction with β_48_ is given in **Supplementary Figure S6**.

The resulting network shown in **Figure 4a** that was built by whitelisting all edges learned in the previous section featured more inter-chain edges than were obtained when learning the network as-is from input data with no constraints. There is, however, little if any difference in the likelihood of TCRαβ complexes compared to the sum of likelihoods from separate chains as shown in **Figure 4b**. Notably, small likelihood values in this figure stemmed from the fact that certain V/J gene combinations are quite rare in the PairSEQ dataset with frequency less than 10^-4^ and that the network did not include all possible edges between intra-chain residues which are in reality all interconnected, leading to the presence of lots of independent probability distribution products.

**Figure 4.**
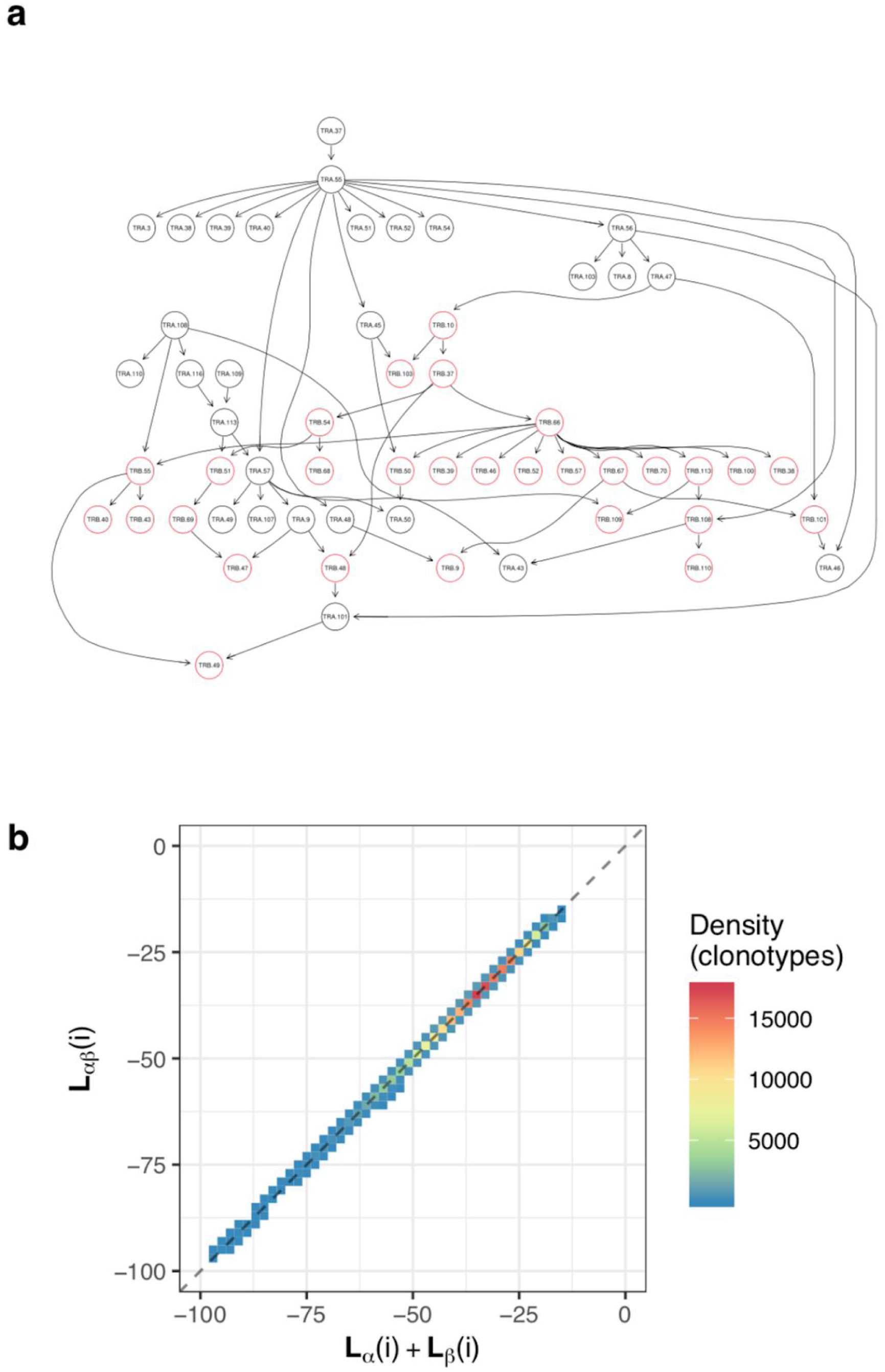
Bayesian network (BN) of TCRαβ complex residues. **a**. The graph of BN built with separately learned inter-chain contacts (shown in **Supplementary Figure S4**) whitelisted and residues that are not contacting according to contact frequency thresholding blacklisted. **b**. A density plot showing correlation between log-likelihood (LL) of BN for paired chains (y axis, computed using the network in **a**.) and sum of LLs of individual α and β chains for i_^th^_clonotype from PairSEQ dataset. In order to compute individual chain LLs two independent networks were built by removing inter-chain edges and separating α and β residue components of the BN.

By computing the entropy based on all allowed amino acid profiles of the selected residues one can observe the following: the entropy of separate α chain is H_α_=4.68, separate β chain is H_β_=2.71, and the entropy of TCRαβ complexes is H_αβ_=7.43. The difference between these values is H_αβ_-H_α_-H_β_=0.04, suggesting that the amount of information conveyed by constraints imposed by α and β chain sequences on chain pairing is negligibly small.

### Amino acids at contact residues influence the conformation of the TCRαβ complex

Given little pairing information encoded in contact residues we suggested that they may still play a role in defining the overall conformation of the TCRαβ complex. We therefore checked mutual orientation of α and β chains of complexes having distinct amino acids at some of the critical contact residues identified in the previous section and shown in **Supplementary Figure S5**. In order to define chain orientation, we first computed principal axes of chain atoms that provide a general description of the characteristics of a rigid body approximating a given chain (**Figure 5a top**), an approach that was previously successfully applied to describe the geometry of both TCR and antibody complexes [13]. We then used these axes to compute Euler angles between α and β chains that provide a comprehensive description of their mutual orientation (**Figure 5a bottom**).

**Figure 5.**
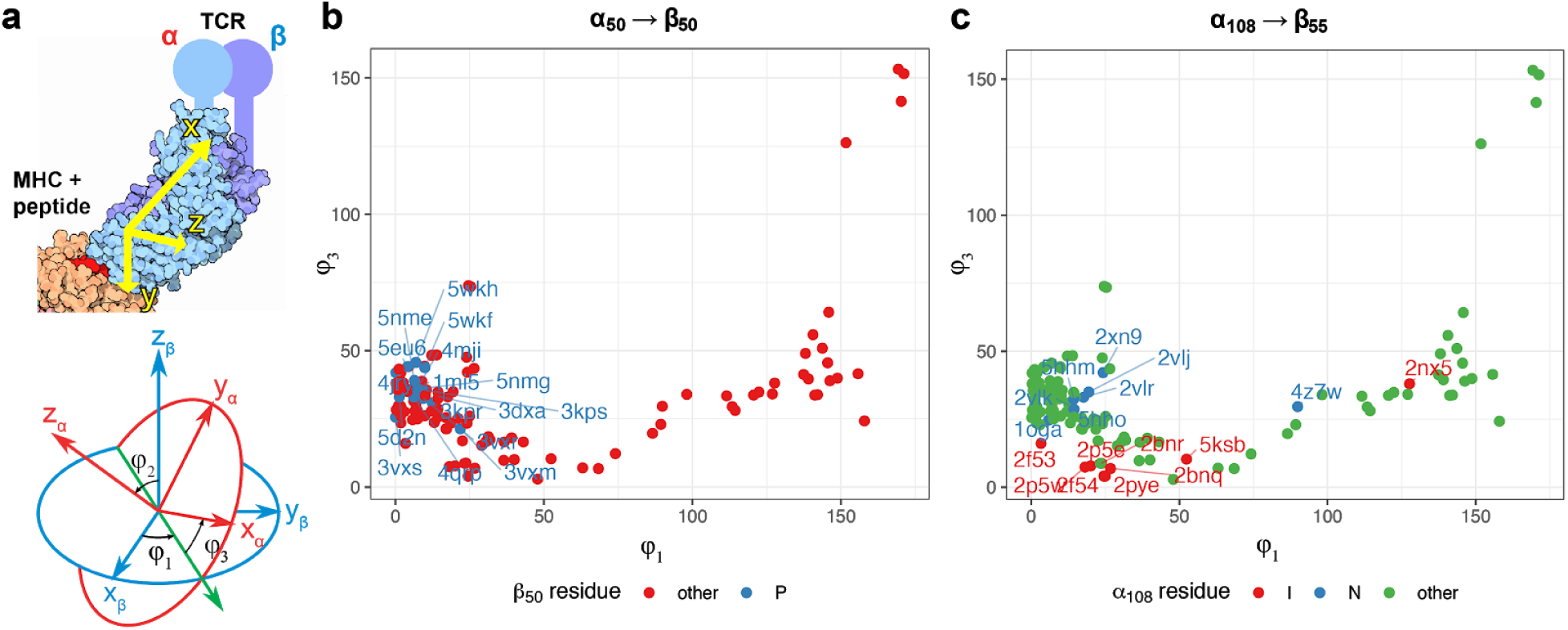
Contacting residues define mutual orientation of chains in TCRαβ complex. **a**. A schematic definition of angles between α and β chains. Principal axes (x_α_, y_α_, z_α_) and (x_β_, y_β_, z_β_) of both TCR chains are computed using the inertia tensor of all atoms of a given chain with the exception of constant domain atoms (**top panel**); representative orientation of principal axes in real TCR:pMHC complex are shown. Euler angles φ_1,2,3_ are then computed by superimposing chain centers of mass and computing angles between α and β principal axes (**bottom panel**). Illustrations were adapted from Wikimedia Commons (https://commons.wikimedia.org/wiki/File:63-T-CellReceptor-MHC.tif and https://commons.wikimedia.org/wiki/File:Eulerangles.svg). **b**. and **c**. - scatter plots of φ_1_ and φ_3_ angles between α and β chains in complexes having α_50_→β_50_ and α_108_→β_55_ contacts respectively (example contacting residue pairs selected from **Supplementary Figure S5**). Note the differences in orientation between chains of TCRαβ complexes: difference in φ_1_ angle between chains for complexes having either Proline (P) or other residue at β_50_ (**b**), and difference in φ_3_ angle for complexes having either Isoleucine (I) or Asparagine (N) at α_108_ (**c**).

To demonstrate that certain amino acid incidence at contacting residues can significantly alter TCRαβ complex conformation we have chosen two representative contacts: α_50_→β_50_ and α_108_→β_55_, and assessed the differences in Euler angles φ_1_ and φ_3_ that can be explained by specific amino acid choice. As shown in **Figure 5b**, the presence of Proline at β_50_ residue substantially restricts the available φ_1_ angle value range: 8° vs 44° on average for Proline versus any other residue (P < 10^-8^, two-tailed T-test). On the other hand, **Figure 5c** demonstrates that the choice of either Isoleucine or Asparagine at α_108_ residue leads to substantially different φ_3_ angle values, 11° and 31° on average respectively (P < 10^-3^, two-tailed T-test). Overall, these examples demonstrated that the choice of specific amino acid at contacting residues can have a significant effect on mutual orientation of TCRαβ complex chains.

### Antigen-driven selection overrides pairing preferences

As overall differences in pairing preferences in TCRαβ complex were relatively subtle, we hypothesized that the TCR repertoire can still show αβ pairing preferences when subject to perturbations, such as antigen-driven selection and expansion. We have therefore investigated pairing preferences in TCRs specific to certain epitopes extracted from the VDJdb database [14] (see **Materials and Methods** section).

To study this case, we computed likelihood values of the observed TCRαβ complexes for the original dataset and datasets where shuffling was performed within and between epitope-specific TCR groups. The likelihoods were computed using a BN that solely included inter-chain contacts (**Supplementary Figure S4**) to minimize the V and J gene usage bias. As demonstrated in **Figure 6**, selection based on epitope preferences can distort pairing preferences, given a rise in both more likely and less likely TCRαβ complexes compared to random pairing. On the other hand, there is little difference in pairing preferences within epitope-specific TCR groups so that shuffling chain pairs doesn’t change the likelihood distribution, and one would expect little constraints on pairing in TCRαβ complexes recognizing the same epitope.

**Figure 6.**
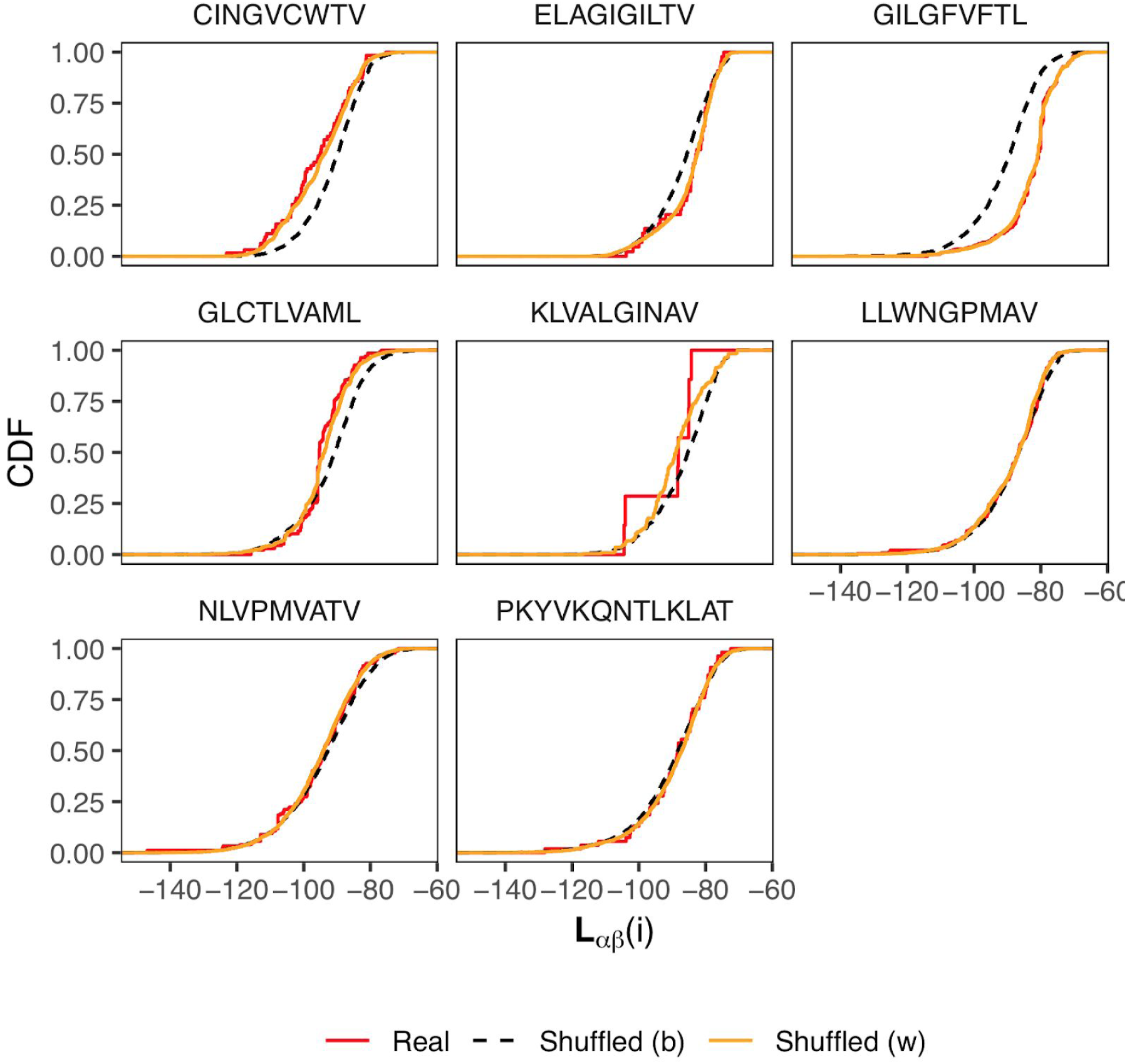
Log-likelihood (LL) distributions for TCRαβ pairs with known antigen specificity. Cumulative distribution functions of αβ pair LLs computed according to the model shown in **Figure 4a**. The plot shows n=1388 real TCRαβ pairs from the VDJdb database (red curves), as well as pairs shuffled within groups specific to the same epitope (orange curves) and pairs shuffled across the entire dataset (dashed black curve). For the latter case, n=10,000 pairs were selected at random for each epitope pair with re-sampling allowed in order to balance the dataset. Labels above panels are the cognate epitope sequences.

### TCRαβ pairing preferences of invariant T-cells

As it is well-known that there are certain subsets of T-cells characterized by invariant T-cell receptor structure such as MAIT [15] and iNKT [16] cells, we decided to investigate selection biases related to T-cell specialization and phenotype. For example, MAIT cells were shown to be enriched in TCR sequences rearranged from TRAV1-2 and TRAJ12/20/33 that are mostly paired with TRBV6-4/20 [17]. As a validation of our framework, we investigated the enrichment of specific αβ contacts characteristic for MAIT cells using the PairSEQ dataset. We compared residue pair frequencies in VαJαVβ of MAIT cells with clones having MAIT VαVβ and any Jα choice (**Figure 7a**). We found several residue pairs that involve Jα and have higher amino acid pair frequency than expected by chance, however, little enrichment for these contacts in the whole dataset suggests that we have found indirect interactions driven by the need to recognize the MR1 molecule by MAIT cells rather than direct αβ contacts.

**Figure 7.**
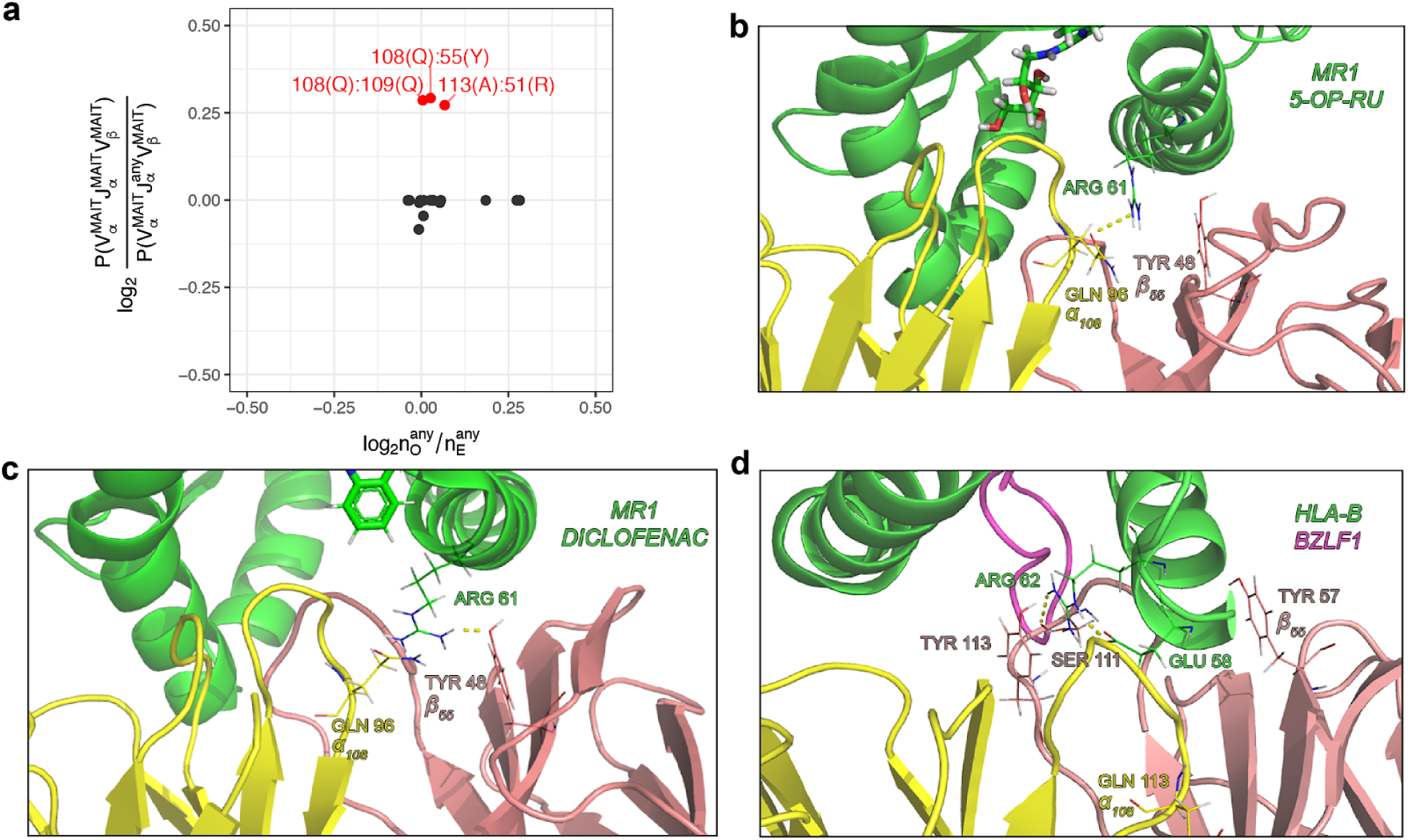
Characteristic residue contacts of MAIT TCRs. **a**. Scatter-plot of amino acid pair enrichment at contacting residues for the Jα gene choice of MAIT T-cells versus overall enrichment observed for given contact residues in the PairSEQ dataset. Y axis shows the log ratio of amino acid pair probabilities for VαJαVβ combinations corresponding to MAIT T-cells and those with a free choice for the Jα gene. X axis shows observed to expected amino acid pair count ratio at contacting residues in the PairSEQ dataset. Residue pairs with enriched amino acid pairs coming from MAIT Jα gene choice (y > 0.25, corresponding to ∼19% increase in frequency) are colored in red and labeled. Not that overall enrichment for corresponding amino acid pairs in the PairSEQ dataset is relatively moderate (x < 0.125, corresponding to ∼9% increase in frequency). **b**.**-d**. Structural data showing Glutamine (GLN, Q) at α_108_ and Tyrosine (TYR, Y) at β_55_ interacting with an Arginine (ARG) of MHC alpha-1 helix domain in the MAIT:MR1 complex. MR1 complex structures are shown in **b** (*4pj7*) and **c** (*5u1r*), **d** shows an non-MAIT TCR (having the same amino acids at α_108_ and β_55_) in complex with MHCI (*4jry*). Polar contacts between GLN:ARG and TYR:ARG are shown with dotted lines in **b** and **c**, but are absent in **d**. PDB structure chain coloring: green for MHC, yellow for TCRα and pink for TCRβ; antigen peptide in **d** and **e** is shown with purple.

We turned to structural data on available MAIT TCR complexes to further investigate contacts enriched in MAIT subset, namely α_108_GLN and β_55_TYR. We visually inspected twenty available MAIT TCR-ligand-MR1 complex structures and a structure of conventional TCR having corresponding residues in Vβ and Jα obtained from the Protein Data Bank and used PyMOL to find contacts involving those residues. While we failed to detect any direct contact between α_108_ and β_55_, we found out that both of them can interact with ARG residue of the alpha-1 helix of the MR1 molecule: MR1 ARG is close to and faces both α_108_GLN and β_55_TYR. In some of MAIT structures (PDB IDs *4pj7, 4pj8* and *5d7j*) MR1 ARG forms a hydrogen bond with α_108_GLN only (**Figure 7b**), while in most of the remaining structures a hydrogen bond with β_55_TYR is formed (**Figure 7c**). At the same time, in the latter case ARG spatial orientation could be stabilized by the α_108_GLN side chain group from below even in the absence of a direct contact, so we suggest that both TCR residues are required to sandwich MR1 ARG. Notably, while there is a similarly placed ARG residue in the alpha-1 helix of a conventional MHCI molecule, it is situated far from corresponding TCR residues and doesn’t form any contacts with them (**Figure 7d**), so the interaction between MR1 ARG, α_108_GLN and β_55_TYR is likely a unique feature of MAIT TCRs.

### De-novo detection of invariant TCRs using αβ pairing biases

Results presented above support the hypothesis of unconstrained chain pairing in TCRαβ complex, and we observe pairing biases in cases of antigen-specific T-cell subsets and MAIT T-cells, but not at the level of entire T-cell repertoire. Thus, we hypothesize that deeper exploration of pairing biases can be a useful instrument for T-cell subset discovery. To demonstrate the feasibility of this method, we analyzed frequencies of VαJαVβJβ gene combinations in the PairSEQ dataset in order to detect cases of biased αβ pairing and explored the potential phenotype of corresponding T-cells using literature and published scRNASeq data. We based our analysis on TCR gene trios, i.e. JαVβJβ, VαVβJβ, VαJαJβ and VαJαVβ combinations. The rationale behind this is that invariant TCRs of MAIT and iNKT cells are commonly defined based on the VαJαVβ gene trio [18]. We next compared the number of times each TCR gene combination is observed in the PairSEQ dataset and its count expected under the assumptions of random αβ pairing using hypergeometric enrichment test (**Figure 8a)**. As can be seen in the figure, conventional MAIT and iNKT cells are clear outliers marking several highly enriched VαJαVβ combinations. Notably, we also observed a high number or enriched TCR gene trios that are of unknown origin and were not described previously (**Supplementary Table 1**).

**Figure 8.**
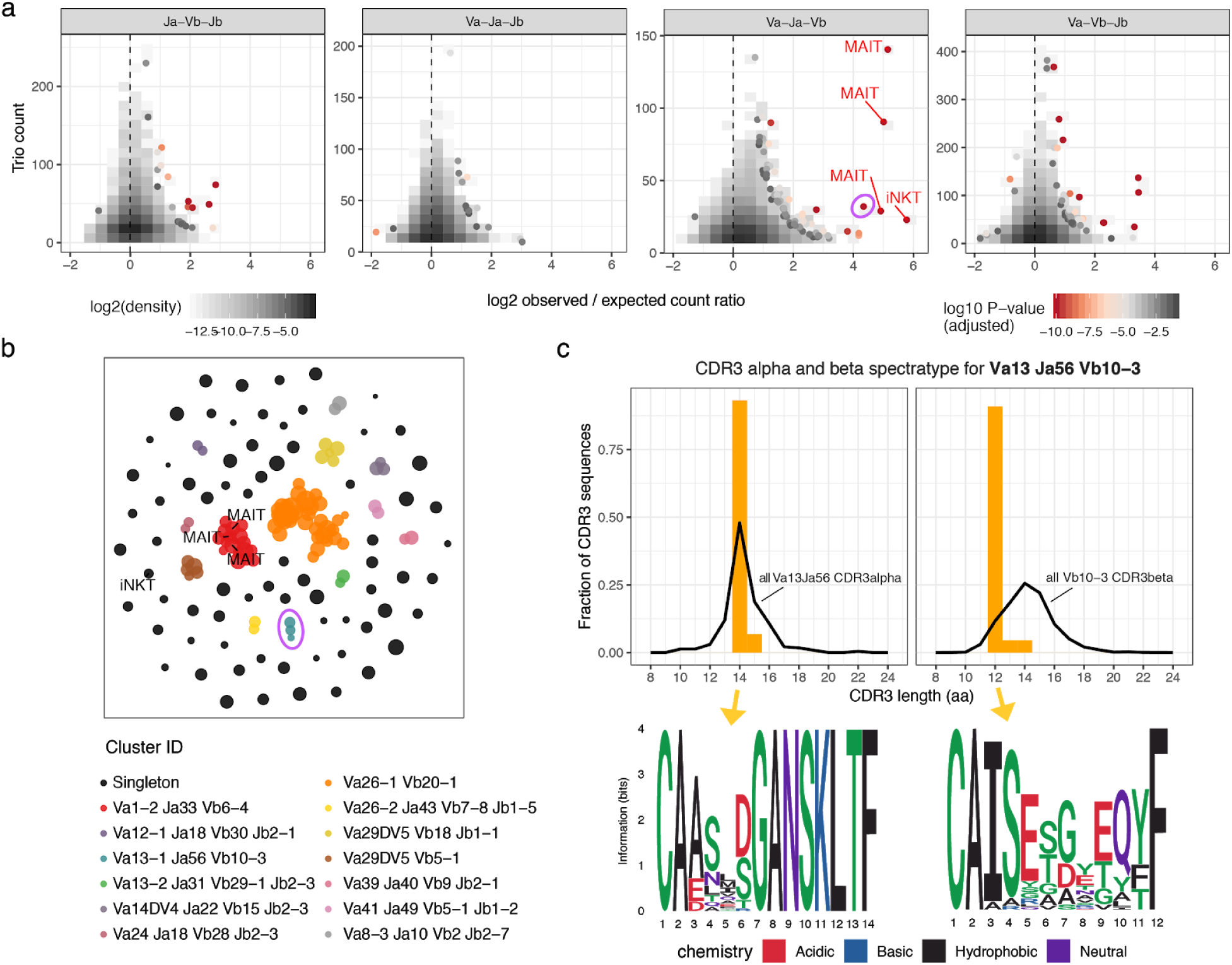
Exploring invariant TCR using enrichment analysis of VαJαVβJβ gene combinations. **a**. Scatterplot showing enrichment of certain TCR gene trios (unique combinations of three of four TCR germline genes, either JαVβJβ, VαVβJβ, VαJαJβ or VαJαVβ) in the PairSEQ dataset. Logarithm of the ratio of the observed and expected counts for all possible gene trios is plotted against their observed count. Expected count is calculated under the assumption of random αβ pairing as (count of α part alone) x (count of β part) / (total number of reads). Points are colored by the P-value of the hypergeometric enrichment test for the co-occurrence of α and β parts of the gene trio (adjusted for multiple comparisons using Holm method). Canonical MAIT (TRAV1-2, TRAJ12/20/33, TRBV6-4) and iNKT (TRAV10, TRAJ18, TRBV25-1) variants are highlighted with corresponding labels. Only gene trios supported by at least 10 reads are shown. Pink circle highlights the Va13 Ja56 Vb10-3 population. **b**. Grouping of selected TCR gene trios (having adjusted P < 0.05 for enrichment test and represented by at least 10 reads) according to overlap between their VαJαVβJβ gene sets. The plot shows the layout of the resulting graph of gene trios (nodes), having edges connecting pairs of nodes with exactly matching gene sets (missing genes, e.g. Vα in JαVβJβ, are considered as wildcards). Nodes of the graph are represented by points and are colored according to the connected component (cluster) of the network they were assigned to. Cluster ID is a combination of most frequent gene names in co-clustered trios. **c**. CDR3 spectratyping and motifs for the Va13 Ja56 Vb10-3 population. Top plots show distribution of CDR3 alpha (left) and beta (right) chains of the population compared to all PairSEQ TCRs rearranged with corresponding alpha or beta segments, note that only a single dominant length is present for both alpha and beta. Bottom plots show sequence logos of corresponding CDR3 lengths in the population.

Grouping enriched TCR gene trios based on partial overlap between gene sets yields a number of large connected components, one of which is clearly linked to MAIT, and a singleton representing iNKT cells (**Figure 8b**). The Va13 Ja56 Vb10-3 population demonstrates a remarkable enrichment (pink circle in **Figure 8ab, Supplementary Table 1**) that is close to MAIT and iNKT cells in magnitude. Further investigation of this population showed that it indeed features invariant TCRs that are dominated by 14aa and 12aa CDR3 alpha and beta sequences encoding a prominent motif (**Figure 8c**).

## Discussion

In the present study we performed a deep and comprehensive analysis of T-cell receptor sequencing and structural data aimed at detecting biases in TCRα and β chain pairing preferences. We were able to identify dozens of TCR residues that are involved in inter-chain contacts, however, even those positions of either chain that can feature several amino acids corresponding to distinct V and J genes were relatively uniform in amino acid pair preference. Most of irregularities we detected were related to cases present in less than 100 cells per 10^5^ T-cells analyzed which, together with information measurement based on inferred residue probability distribution, provides a solid proof for nearly random pairing of TCRα and β chains. The consequence of our finding is that human T-cell repertoire in general feature almost random combination of TCR α and β chains. This observation is in line with one of the aims of the adaptive immune system, which is to field the most diverse repertoire of T-cells in order to be able to successfully combat newly encountered pathogens.

One of the limitations of analysis is that it was aimed at germline parts of TCR as it is almost impossible to summarize the impact of the central part of CDR3 region that is mostly involved in antigen recognition and is hard to bring to a uniform indexing due to a huge variety of potential conformations of the resulting loop. We checked potential associations between various CDR3 features and failed to find any substantial correlations between values for alpha and beta chain CDR3 regions (**Supplementary Figure 7**).

Specific choice of contacting amino acids can nevertheless influence the overall conformation of the TCRαβ complex. As mutual orientation of chains affected by the choice of specific germline-encoded residues of V and J genes, knowing the possible spectrum of residues that affect conformation and corresponding orientations can aid in de-novo TCR:peptide:MHC complex reconstruction [19]. As only a handful of Vα/Jα/Vβ/Jβ allele combinations are currently present in the set of PDB structures, predictive models aimed at specific contacting residues can greatly extend the number of templates available for this task.

As expected, pairing in TCRαβ complexes can be heavily distorted by T-cell selection. Our results suggest that the selection of TCRs based on antigen specificity can lead to both more and less frequent αβ pairs depending on the antigen, suggesting that antigenic stimuli can override the uniform landscape of αβ pairing. Notably, chain pairing within the T-cell subset specific to the same epitope appears to be almost random, which is in line with our previous observations [20] and suggests that there is little competition between α and β chains in epitope recognition.

Our observation of random pairing in TCRαβ complexes in the absence of T-cell selection and the fact that there are specific T-cell subsets characterized by invariant TCR sequences lead us to explore pairing preferences in these subsets. Our results show that there is no intrinsic bias for interaction between α and β chains in invariant MAIT cells: observed pairing preferences stem from the need of a cooperative interaction between TCR α and β chains and the MR1 molecule recognized by MAIT cells. Furthermore, we were able to link observed irregularities in α and β chain pairing to a previously undescribed invariant TCR, suggesting that our results can serve as a basis for detection of novel specialized T-cell subsets.

There are some open questions not discussed in the study, such as whether inter-chain interactions in TCRαβ complex can lead to distinct configurations that are more or less preferable for targeting specific epitopes. The methodology in this manuscript is can be also extended to account for interaction between germline parts of the TCRαβ and MHC molecules, potentially leading to models describing docking and binding preferences in the TCRαβ:MHC complex.

## Acknowledgements

The authors would like to thank Ivan Zvyagin, Vadim Karnaukhov, Pavel V Shelyakin, Alexey Davydov and Maria Metsger for fruitful discussions. This study was supported by Russian Science Foundation Grant No. 17-15-01495.

## Supplementary Figures and Tables

**Supplementary Figure S1.**
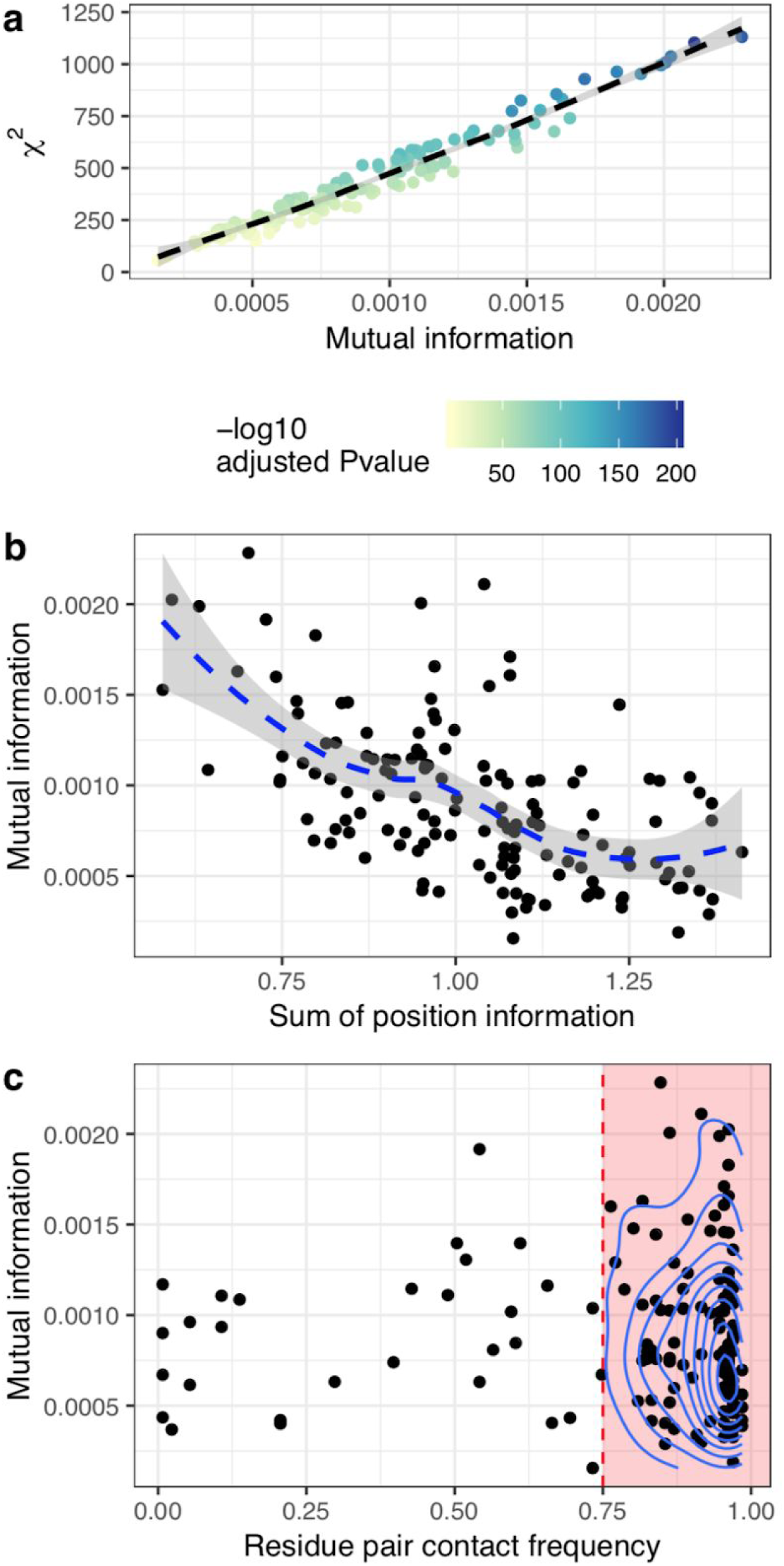
Amino acid pairing statistics at contacting residues. **A**. Scatter-plot of *χ*^2^ statistic values versus mutual information for AA pair frequency matrices of each contacting residue pair. **b**. Mutual information is negatively correlated with conservation (information content) of V and J germline residues. **c**. Scatterplot of residue pair contact frequencies and mutual information of AA pair frequency matrices.

**Supplementary Figure S2.**
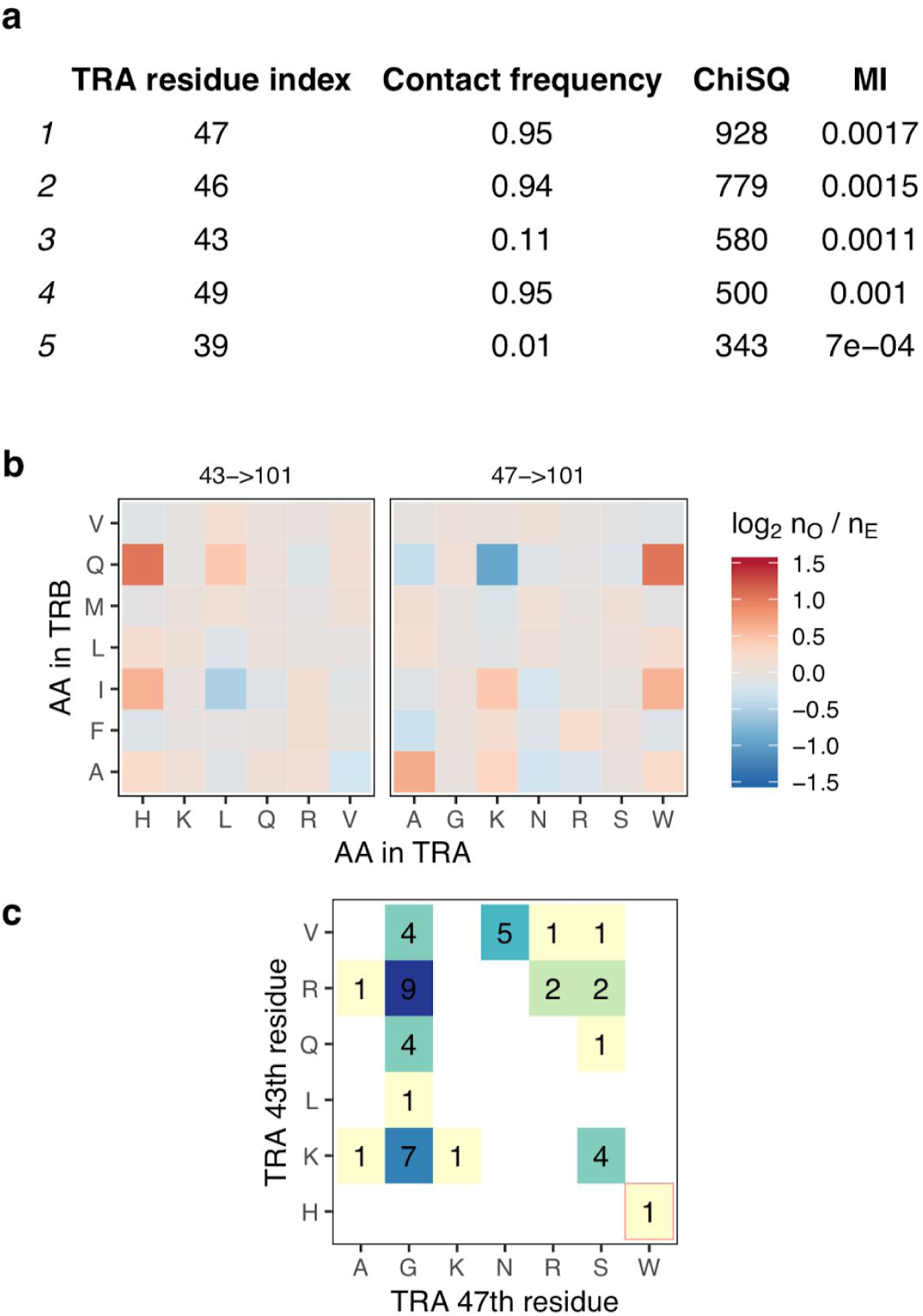
A case of an indirect correlation between residue amino acid profiles. **a**. Contact frequency, *χ*^2^ and mutual information for various TCRα residues contacting β_101_. Note that α_47_ is contacted ∼95% of times, while α_43_ is only contacted at a rate of ∼11%. **b**. Tables of observed to expected contact count ratios showing highly non-random profiles for Histidine in α_43_ and Tryptophan in α_47_. The correlation between these two table columns is R ≈ 1. **c**. Co-occurrence of amino acids between α_43_ and α_47_ in the set of human V genes. Numbers indicates the quantity of V genes with a given combination. The gene having Histidine and Tryptophan in corresponding positions is highlighted with a red box.

**Supplementary Figure S3.**
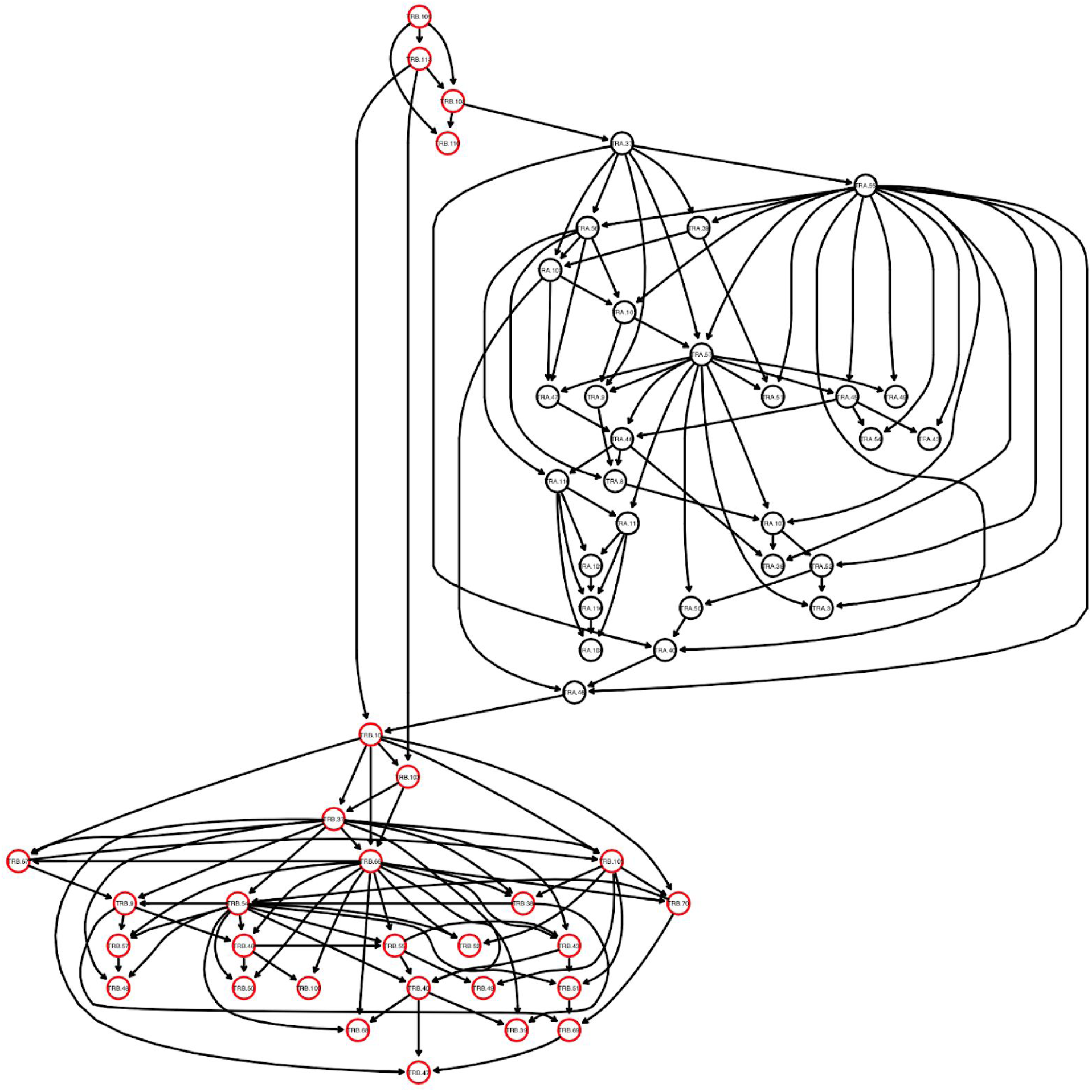
Bayes network of residue amino acid profiles. The plot shows a graph encoding dependencies between amino acid profiles of all TCRαβ complex residues built for the entire PairSEQ dataset with no constraints on allowed and disallowed edges. Black and red nodes show TCRα and TCRβ residues respectively. The only inter-chain contacts inferred are β_108_→α_37_, and α_46_→β_10_.

**Supplementary Figure S4.**
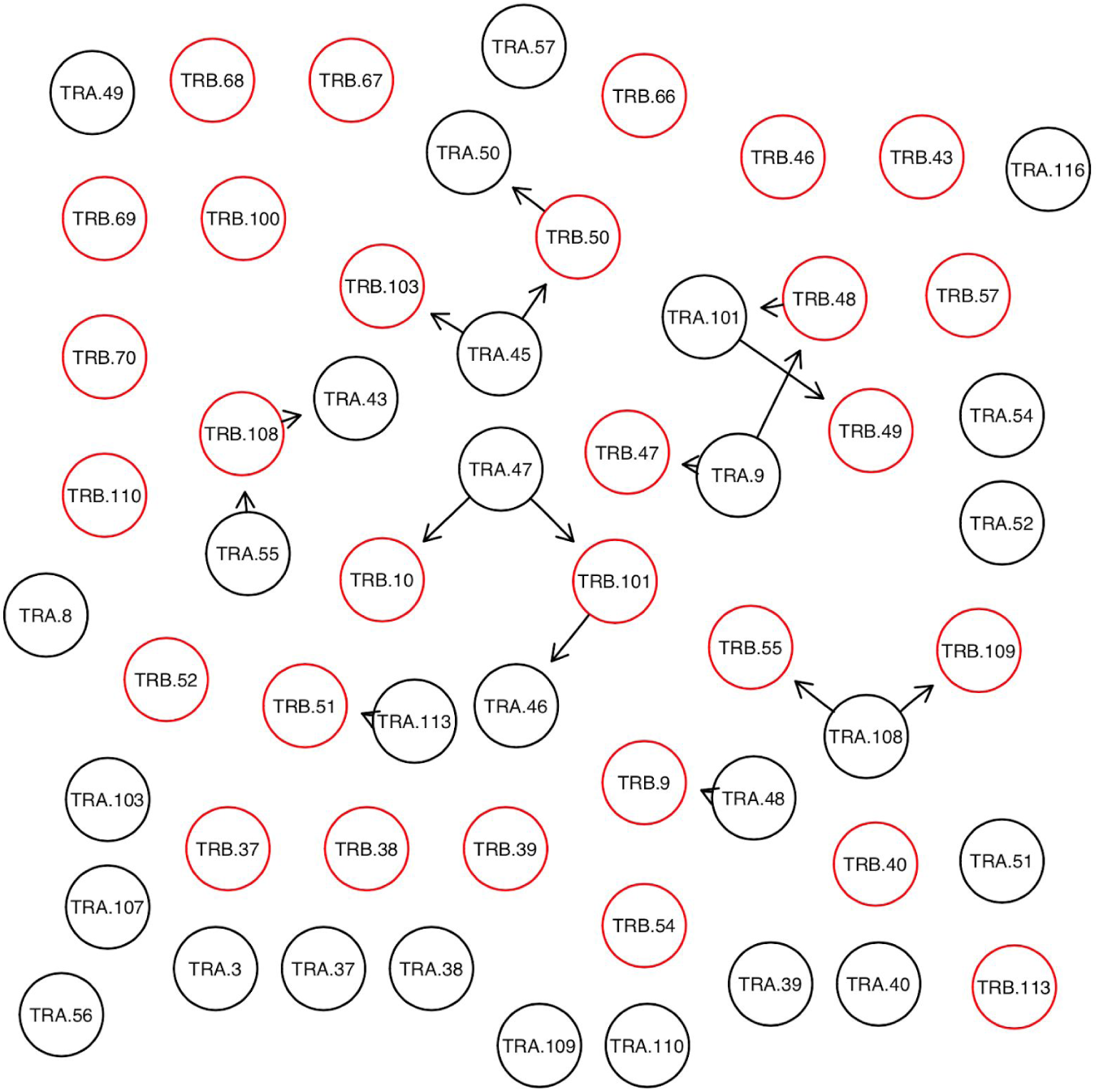
Bayes network of inter-chain contacts. The plot shows a graph encoding dependencies between amino acid profiles of selected contacting residues built for the entire PairSEQ dataset with only inter-chain residue edges allowed. Black and red nodes show TCRα and TCRβ residues respectively.

**Supplementary Figure S5.**
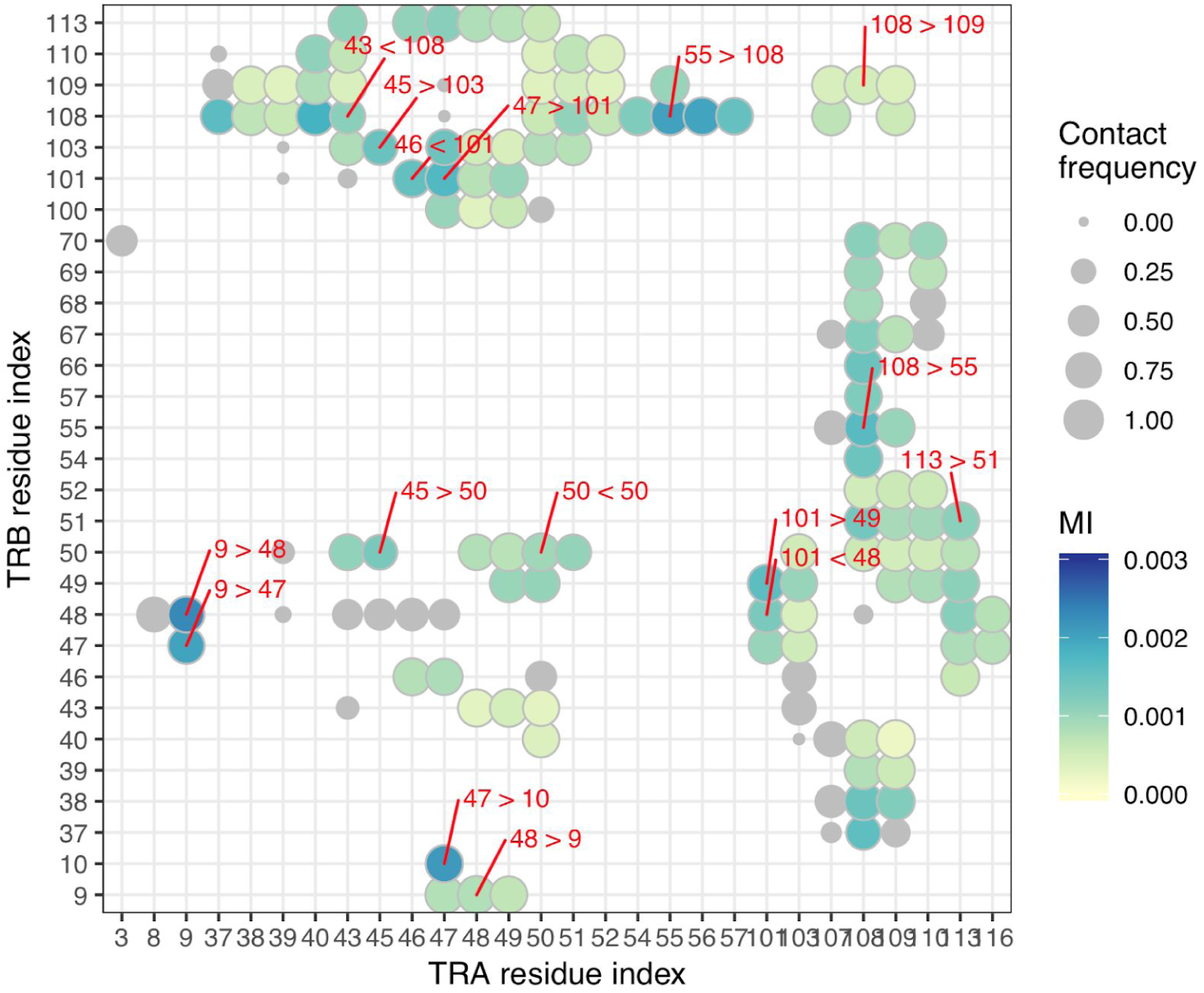
Contact map with inferred direct αβ contacts. A contact map of TCRαβ complexes with contacting residue pairs inferred by Bayes network (BN) analysis shown in **Supplementary Figure S4** highlighted with labels. Greater (>) and less (<) signs show node directionality in BN, first and second numbers in node label represent α and β chain residue indices respectively. Node color shows mutual information of amino acid pair frequency matrices. Node size represents contact frequency for residue pairs inferred from structural data, grey nodes are the ones that fall below 0.75 contact frequency threshold.

**Supplementary Figure S6.**
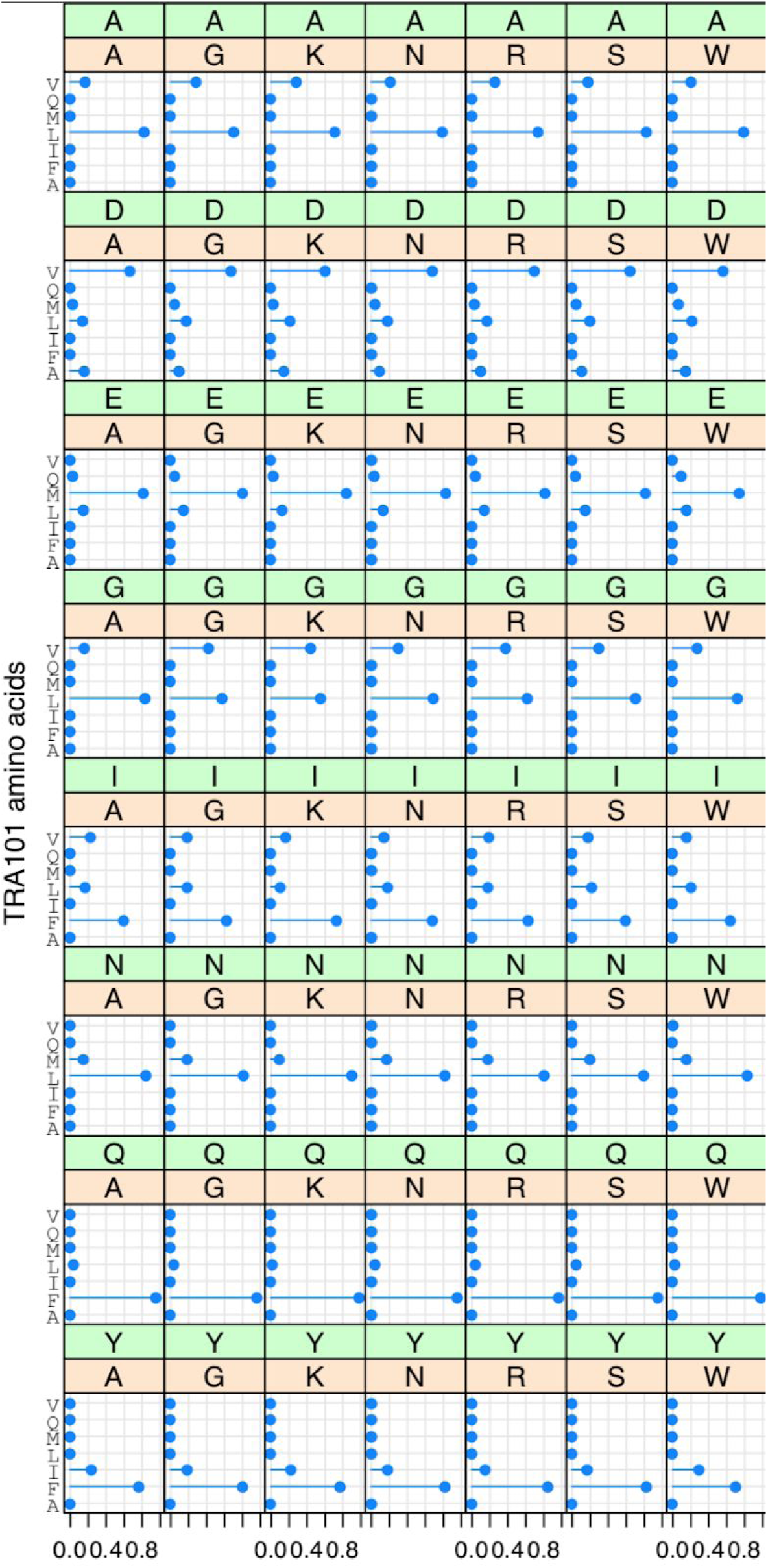
Representative conditional probabilities learned using Bayes network. Conditional probabilities for residue α_101_ given residue α_55_ (green labels above panels) and residue β_48_ (orange labels). Subtle differences for e.g. Leucine (L) at α_101_ between Glycine (G) and Lysine (K) at β_48_ given Isoleucine (I) at α_55_ can be noted.

**Supplementary Figure 7.**
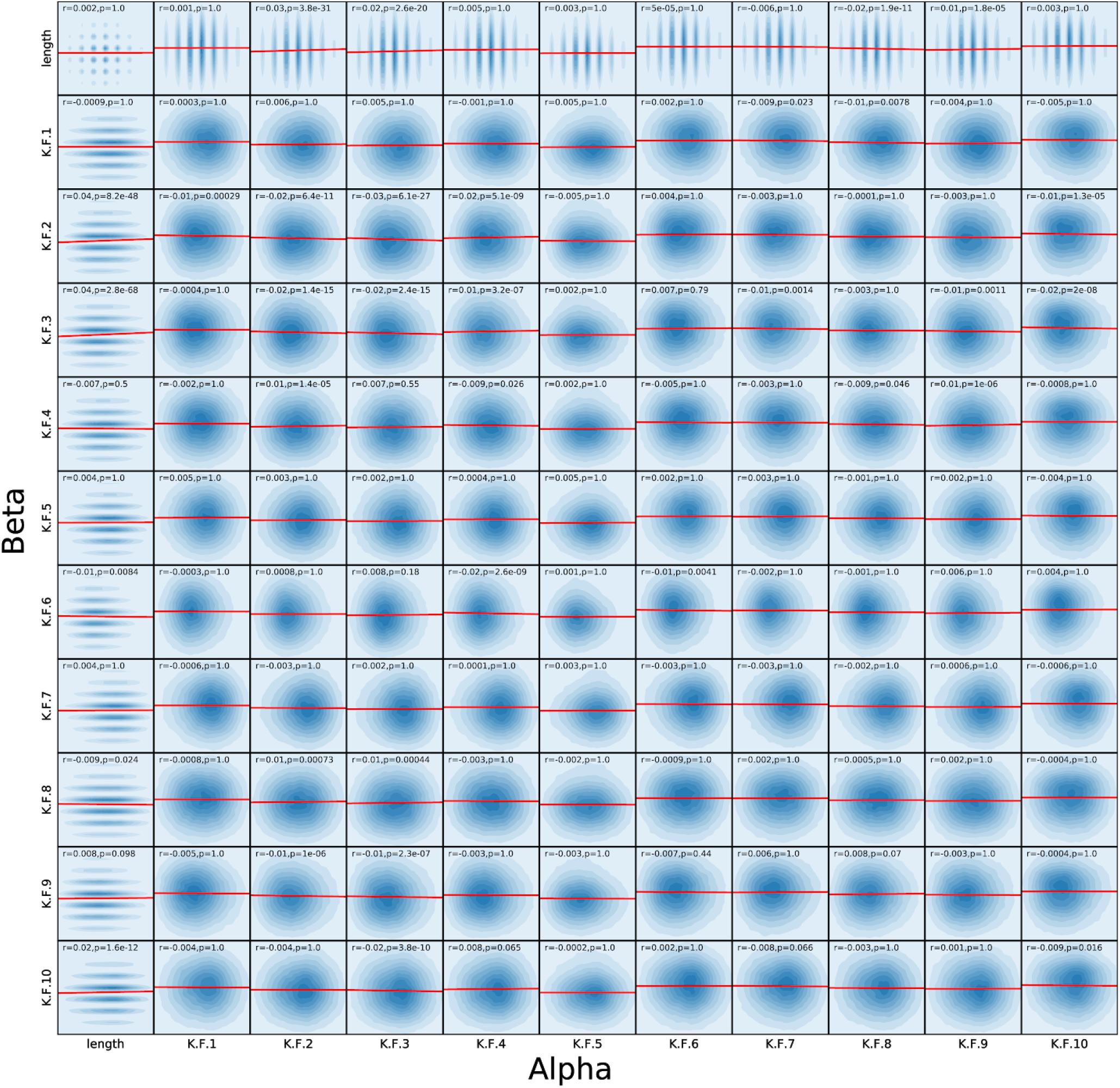
Correlation analysis of CDR3 features of paired TCR alpha and beta chains. CDR3 length and means of Kidera factors (KF1-10) are shown. Pearson correlation coefficients between alpha and beta chain values and corresponding Bonferroni-adjusted P-values are provided above each subplot.

**Supplementary Table 1.** Table of enriched V/J gene trios. Cluster ID specifies groups of trios that have overlapping genes. Known subset specifies canonical MAIT or iNKT rearrangements. Observed (obs) and expected (exp, given random alpha-beta pairing) counts are shown for the PairSEQ dataset. A multiple testing-adjusted P-value of a hypergeometric enrichment test is provided for each gene trio.

| Cluster ID | Va | Ja | Vb | Jb | Known subset | Count (obs) | Count (exp) | Adjusted P-value |
| --- | --- | --- | --- | --- | --- | --- | --- | --- |
| Va8-3 Ja10 Vb2 Jb2-7 | X | TRAJ10 | TRBV2 | TRBJ2-7 |  | 85 | 35 | 4.3E-08 |
| Va8-3 Ja10 Vb2 Jb2-7 | TRAV8-3 | TRAJ10 | TRBV2 | X |  | 45 | 15 | 1.4E-05 |
| Va41 Ja49 Vb5-1 Jb1-2 | TRAV41 | TRAJ49 | TRBV5-1 | X |  | 60 | 26 | 7.3E-05 |
| Va41 Ja49 Vb5-1 Jb1-2 | TRAV41 | X | TRBV5-1 | TRBJ1-2 |  | 43 | 18 | 1.2E-02 |
| Va39 Ja40 Vb9 Jb2-1 | TRAV39 | X | TRBV9 | TRBJ2-1 |  | 72 | 32 | 3.7E-05 |
| Va39 Ja40 Vb9 Jb2-1 | TRAV39 | TRAJ40 | TRBV9 | X |  | 31 | 10 | 4.4E-04 |
| Va29DV5 Vb5-1 | TRAV29DV5 | TRAJ39 | TRBV5-1 | X |  | 56 | 21 | 4.3E-06 |
| Va29DV5 Vb5-1 | TRAV29DV5 | TRAJ6 | TRBV5-1 | X |  | 22 | 4 | 5.1E-06 |
| Va29DV5 Vb5-1 | TRAV29DV5 | X | TRBV5-1 | TRBJ1-5 |  | 55 | 24 | 7.6E-04 |
| Va29DV5 Vb5-1 | TRAV29DV5 | X | TRBV5-1 | TRBJ2-1 |  | 168 | 110 | 1.1E-02 |
| Va29DV5 Vb5-1 | TRAV29DV5 | TRAJ52 | TRBV5-1 | X |  | 52 | 24 | 1.7E-02 |
| Va29DV5 Vb18 Jb1-1 | TRAV29DV5 | X | TRBV18 | TRBJ1-5 |  | 66 | 26 | 5.1E-07 |
| Va29DV5 Vb18 Jb1-1 | TRAV29DV5 | X | TRBV18 | TRBJ1-1 |  | 89 | 44 | 3.3E-05 |
| Va29DV5 Vb18 Jb1-1 | TRAV29DV5 | X | TRBV18 | TRBJ1-2 |  | 70 | 31 | 1.0E-04 |
| Va29DV5 Vb18 Jb1-1 | TRAV29DV5 | TRAJ24 | TRBV18 | X |  | 19 | 5 | 5.8E-03 |
| Va29DV5 Vb18 Jb1-1 | TRAV29DV5 | X | TRBV18 | TRBJ1-6 |  | 40 | 17 | 1.9E-02 |
| Va26-2 Ja43 Vb7-8 Jb1-5 | TRAV26-2 | X | TRBV7-8 | TRBJ1-5 |  | 38 | 13 | 1.5E-03 |
| Va26-2 Ja43 Vb7-8 Jb1-5 | TRAV26-2 | TRAJ43 | TRBV7-8 | X |  | 27 | 8 | 2.6E-03 |
| Va26-1 Vb20-1 | TRAV12-3 | X | TRBV20-1 | TRBJ2-3 |  | 260 | 149 | 0.0E+00 |
| Va26-1 Vb20-1 | TRAV26-1 | X | TRBV20-1 | TRBJ1-1 |  | 216 | 113 | 0.0E+00 |
| Va26-1 Vb20-1 | TRAV26-1 | X | TRBV20-1 | TRBJ1-3 |  | 97 | 35 | 0.0E+00 |
| Va26-1 Vb20-1 | TRAV26-1 | X | TRBV20-1 | TRBJ2-1 |  | 368 | 237 | 1.6E-11 |
| Va26-1 Vb20-1 | TRAV13-1 | TRAJ17 | TRBV20-1 | X |  | 90 | 38 | 1.6E-10 |
| Va26-1 Vb20-1 | X | TRAJ10 | TRBV20-1 | TRBJ1-4 |  | 46 | 12 | 9.6E-10 |
| Va26-1 Vb20-1 | X | TRAJ17 | TRBV20-1 | TRBJ1-2 |  | 122 | 58 | 6.4E-09 |
| Va26-1 Vb20-1 | TRAV12-3 | TRAJ18 | TRBV6-4 | X |  | 14 | 1 | 9.5E-09 |
| Va26-1 Vb20-1 | TRAV26-1 | X | TRBV20-1 | TRBJ1-2 |  | 199 | 118 | 1.4E-07 |
| Va26-1 Vb20-1 | TRAV12-3 | TRAJ17 | TRBV20-1 | X |  | 75 | 33 | 5.7E-07 |
| Va26-1 Vb20-1 | X | TRAJ3 | TRBV20-1 | TRBJ1-1 |  | 99 | 48 | 4.5E-06 |
| Va26-1 Vb20-1 | TRAV26-1 | TRAJ49 | TRBV20-1 | X |  | 94 | 48 | 1.1E-05 |
| Va26-1 Vb20-1 | X | TRAJ29 | TRBV20-1 | TRBJ1-1 |  | 116 | 61 | 1.3E-05 |
| Va26-1 Vb20-1 | TRAV12-3 | TRAJ48 | TRBV20-1 | X |  | 62 | 26 | 1.5E-05 |
| Va26-1 Vb20-1 | TRAV9-2 | TRAJ23 | TRBV20-1 | X |  | 98 | 52 | 2.6E-05 |
| Va26-1 Vb20-1 | TRAV26-1 | TRAJ23 | TRBV20-1 | X |  | 98 | 52 | 4.2E-05 |
| Va26-1 Vb20-1 | TRAV12-3 | X | TRBV20-1 | TRBJ2-5 |  | 200 | 128 | 7.3E-05 |
| Va26-1 Vb20-1 | TRAV12-3 | TRAJ41 | TRBV20-1 | X |  | 38 | 14 | 1.0E-04 |
| Va26-1 Vb20-1 | TRAV12-3 | TRAJ10 | TRBV20-1 | X |  | 77 | 38 | 1.4E-04 |
| Va26-1 Vb20-1 | TRAV12-3 | TRAJ23 | TRBV20-1 | X |  | 79 | 40 | 2.9E-04 |
| Va26-1 Vb20-1 | TRAV4 | TRAJ15 | TRBV20-1 | X |  | 40 | 15 | 4.9E-04 |
| Va26-1 Vb20-1 | TRAV6 | TRAJ23 | TRBV20-1 | X |  | 58 | 27 | 6.3E-04 |
| Va26-1 Vb20-1 | TRAV26-1 | TRAJ30 | TRBV20-1 | X |  | 70 | 35 | 1.2E-03 |
| Va26-1 Vb20-1 | X | TRAJ24 | TRBV20-1 | TRBJ2-5 |  | 94 | 50 | 1.2E-03 |
| Va26-1 Vb20-1 | TRAV13-1 | X | TRBV20-1 | TRBJ2-1 |  | 382 | 288 | 2.4E-03 |
| Va26-1 Vb20-1 | X | TRAJ23 | TRBV20-1 | TRBJ2-1 |  | 230 | 159 | 2.8E-03 |
| Va26-1 Vb20-1 | TRAV26-1 | TRAJ29 | TRBV20-1 | X |  | 80 | 43 | 4.6E-03 |
| Va26-1 Vb20-1 | TRAV26-1 | X | TRBV20-1 | TRBJ2-7 |  | 365 | 276 | 7.0E-03 |
| Va26-1 Vb20-1 | TRAV26-1 | TRAJ39 | TRBV20-1 | X |  | 92 | 53 | 7.2E-03 |
| Va26-1 Vb20-1 | TRAV12-3 | TRAJ18 | X | TRBJ2-3 |  | 40 | 17 | 7.8E-03 |
| Va26-1 Vb20-1 | TRAV26-1 | TRAJ43 | TRBV20-1 | X |  | 77 | 42 | 1.1E-02 |
| Va26-1 Vb20-1 | X | TRAJ15 | TRBV20-1 | TRBJ2-1 |  | 161 | 106 | 2.0E-02 |
| Va26-1 Vb20-1 | TRAV12-3 | TRAJ11 | TRBV20-1 | X |  | 55 | 27 | 2.9E-02 |
| Va26-1 Vb20-1 | TRAV26-1 | TRAJ41 | TRBV20-1 | X |  | 57 | 29 | 3.0E-02 |
| Va26-1 Vb20-1 | TRAV26-1 | TRAJ34 | TRBV20-1 | X |  | 75 | 41 | 4.2E-02 |
| Va26-1 Vb20-1 | TRAV1-1 | TRAJ24 | X | TRBJ2-5 |  | 38 | 16 | 4.6E-02 |
| Va24 Ja18 Vb28 Jb2-3 | TRAV24 | TRAJ18 | TRBV28 | X |  | 21 | 4 | 2.1E-05 |
| Va24 Ja18 Vb28 Jb2-3 | TRAV24 | TRAJ18 | X | TRBJ2-3 |  | 25 | 9 | 3.1E-02 |
| Va14DV4 Ja22 Vb15 Jb2-3 | TRAV14DV4 | TRAJ22 | TRBV15 | X |  | 30 | 4 | 0.0E+00 |
| Va14DV4 Ja22 Vb15 Jb2-3 | TRAV14DV4 | X | TRBV15 | TRBJ2-3 |  | 52 | 17 | 1.1E-07 |
| Va14DV4 Ja22 Vb15 Jb2-3 | TRAV14DV4 | TRAJ22 | X | TRBJ2-3 |  | 73 | 32 | 6.7E-07 |
| Va13-2 Ja31 Vb29-1 Jb2-3 | TRAV13-2 | TRAJ31 | TRBV29-1 | X |  | 25 | 6 | 2.2E-05 |
| Va13-2 Ja31 Vb29-1 Jb2-3 | TRAV13-2 | X | TRBV29-1 | TRBJ2-3 |  | 59 | 29 | 3.6E-02 |
| Va13-1 Ja56 Vb10-3 | TRAV13-1 | TRAJ56 | TRBV10-3 | X |  | 32 | 2 | 0.0E+00 |
| Va13-1 Ja56 Vb10-3 | TRAV13-2 | TRAJ56 | TRBV10-3 | X |  | 12 | 1 | 1.8E-04 |
| Va13-1 Ja56 Vb10-3 | X | TRAJ56 | TRBV10-3 | TRBJ2-7 |  | 24 | 7 | 2.7E-02 |
| Va12-1 Ja18 Vb30 Jb2-1 | TRAV12-1 | TRAJ18 | TRBV30 | X |  | 22 | 4 | 9.5E-05 |
| Va12-1 Ja18 Vb30 Jb2-1 | X | TRAJ18 | TRBV30 | TRBJ2-1 |  | 27 | 9 | 3.2E-02 |
| Va1-2 Ja33 Vb6-4 | TRAV1-2 | TRAJ12 | TRBV6-4 | X | MAIT | 29 | 1 | 0.0E+00 |
| Va1-2 Ja33 Vb6-4 | TRAV1-2 | TRAJ20 | TRBV6-4 | X | MAIT | 91 | 3 | 0.0E+00 |
| Va1-2 Ja33 Vb6-4 | TRAV1-2 | TRAJ33 | TRBV6-4 | X | MAIT | 141 | 4 | 0.0E+00 |
| Va1-2 Ja33 Vb6-4 | TRAV1-2 | X | TRBV6-4 | TRBJ2-1 |  | 138 | 13 | 0.0E+00 |
| Va1-2 Ja33 Vb6-4 | TRAV1-2 | X | TRBV6-4 | TRBJ2-2 |  | 35 | 3 | 0.0E+00 |
| Va1-2 Ja33 Vb6-4 | TRAV1-2 | X | TRBV6-4 | TRBJ2-3 |  | 107 | 10 | 0.0E+00 |
| Va1-2 Ja33 Vb6-4 | X | TRAJ20 | TRBV6-4 | TRBJ2-1 |  | 53 | 14 | 0.0E+00 |
| Va1-2 Ja33 Vb6-4 | X | TRAJ33 | TRBV6-4 | TRBJ2-1 |  | 74 | 10 | 0.0E+00 |
| Va1-2 Ja33 Vb6-4 | X | TRAJ33 | TRBV6-4 | TRBJ2-3 |  | 49 | 8 | 0.0E+00 |
| Va1-2 Ja33 Vb6-4 | X | TRAJ20 | TRBV6-4 | TRBJ2-3 |  | 45 | 11 | 3.3E-11 |
| Va1-2 Ja33 Vb6-4 | TRAV1-2 | TRAJ41 | TRBV6-4 | X |  | 12 | 1 | 3.1E-08 |
| Va1-2 Ja33 Vb6-4 | X | TRAJ33 | TRBV6-4 | TRBJ2-2 |  | 19 | 3 | 1.4E-06 |
| Va1-2 Ja33 Vb6-4 | TRAV1-2 | TRAJ33 | X | TRBJ2-1 |  | 194 | 125 | 2.3E-05 |
| Va1-2 Ja33 Vb6-4 | TRAV1-2 | X | TRBV6-4 | TRBJ2-6 |  | 12 | 1 | 2.9E-05 |
| Va1-2 Ja33 Vb6-4 | TRAV1-2 | TRAJ33 | X | TRBJ2-6 |  | 43 | 15 | 7.4E-05 |
| Va1-2 Ja33 Vb6-4 | TRAV1-2 | TRAJ33 | TRBV4-2 | X |  | 41 | 14 | 1.5E-04 |
| Va1-2 Ja33 Vb6-4 | TRAV1-2 | TRAJ33 | TRBV20-1 | X |  | 135 | 82 | 3.9E-04 |
| Va1-2 Ja33 Vb6-4 | TRAV1-2 | TRAJ33 | TRBV4-3 | X |  | 44 | 19 | 1.8E-02 |
| Va1-2 Ja33 Vb6-4 | TRAV1-2 | X | TRBV4-2 | TRBJ2-1 |  | 51 | 24 | 4.0E-02 |
| Singleton | TRAV10 | TRAJ18 | TRBV25-1 | X | iNKT | 23 | 0 | 0.0E+00 |
| Singleton | TRAV20 | X | TRBV6-2 | TRBJ1-1 |  | 44 | 9 | 0.0E+00 |
| Singleton | TRAV20 | X | TRBV6-3 | TRBJ1-1 |  | 44 | 9 | 0.0E+00 |
| Singleton | TRAV41 | TRAJ50 | TRBV2 | X |  | 15 | 1 | 1.6E-10 |
| Singleton | TRAV21 | X | TRBV27 | TRBJ2-3 |  | 104 | 46 | 2.5E-09 |
| Singleton | TRAV8-1 | TRAJ34 | TRBV28 | X |  | 37 | 10 | 2.8E-07 |
| Singleton | TRAV8-3 | TRAJ34 | TRBV15 | X |  | 27 | 5 | 5.2E-07 |
| Singleton | TRAV22 | X | TRBV29-1 | TRBJ2-7 |  | 50 | 17 | 1.4E-06 |
| Singleton | TRAV20 | TRAJ22 | TRBV30 | X |  | 25 | 5 | 2.0E-06 |
| Singleton | TRAV38-2DV8 | TRAJ9 | TRBV7-9 | X |  | 15 | 2 | 3.4E-06 |
| Singleton | TRAV12-1 | TRAJ48 | TRBV24-1 | X |  | 20 | 3 | 4.6E-06 |
| Singleton | TRAV26-2 | X | TRBV27 | TRBJ1-1 |  | 50 | 18 | 1.9E-05 |
| Singleton | TRAV1-1 | X | TRBV28 | TRBJ2-7 |  | 101 | 52 | 4.3E-05 |
| Singleton | TRAV13-2 | TRAJ6 | TRBV14 | X |  | 20 | 4 | 4.4E-05 |
| Singleton | TRAV34 | TRAJ26 | X | TRBJ1-6 |  | 13 | 2 | 5.9E-05 |
| Singleton | TRAV6 | TRAJ16 | TRBV20-1 | X |  | 40 | 15 | 7.2E-05 |
| Singleton | TRAV14DV4 | TRAJ28 | TRBV9 | X |  | 33 | 10 | 7.7E-05 |
| Singleton | TRAV39 | TRAJ45 | TRBV20-1 | X |  | 61 | 28 | 1.7E-04 |
| Singleton | TRAV26-1 | TRAJ28 | TRBV9 | X |  | 36 | 12 | 3.3E-04 |
| Singleton | X | TRAJ4 | TRBV29-1 | TRBJ2-5 |  | 46 | 17 | 4.8E-04 |
| Singleton | TRAV8-3 | TRAJ21 | TRBV14 | X |  | 14 | 2 | 5.1E-04 |
| Singleton | TRAV3 | TRAJ42 | TRBV30 | X |  | 22 | 5 | 7.3E-04 |
| Singleton | TRAV25 | X | TRBV19 | TRBJ1-5 |  | 65 | 30 | 7.8E-04 |
| Singleton | TRAV12-1 | TRAJ17 | TRBV2 | X |  | 30 | 9 | 9.2E-04 |
| Singleton | TRAV2 | TRAJ15 | TRBV2 | X |  | 31 | 10 | 1.0E-03 |
| Singleton | TRAV5 | TRAJ34 | TRBV28 | X |  | 35 | 12 | 1.1E-03 |
| Singleton | TRAV35 | TRAJ42 | X | TRBJ1-2 |  | 68 | 34 | 1.1E-03 |
| Singleton | TRAV25 | TRAJ11 | TRBV7-9 | X |  | 18 | 4 | 1.5E-03 |
| Singleton | TRAV8-3 | TRAJ8 | TRBV28 | X |  | 36 | 13 | 1.6E-03 |
| Singleton | TRAV12-1 | TRAJ43 | X | TRBJ1-1 |  | 89 | 48 | 1.8E-03 |
| Singleton | TRAV29DV5 | X | TRBV7-9 | TRBJ1-2 |  | 89 | 48 | 2.0E-03 |
| Singleton | TRAV13-2 | TRAJ18 | TRBV24-1 | X |  | 10 | 1 | 2.1E-03 |
| Singleton | TRAV13-2 | X | TRBV7-2 | TRBJ1-5 |  | 57 | 26 | 2.3E-03 |
| Singleton | TRAV8-3 | TRAJ49 | X | TRBJ2-4 |  | 24 | 7 | 2.6E-03 |
| Singleton | TRAV9-2 | TRAJ57 | TRBV12-5 | X |  | 12 | 2 | 3.5E-03 |
| Singleton | TRAV8-1 | TRAJ6 | TRBV30 | X |  | 20 | 5 | 3.7E-03 |
| Singleton | TRAV10 | TRAJ12 | TRBV7-9 | X |  | 13 | 2 | 4.3E-03 |
| Singleton | TRAV13-2 | X | TRBV7-8 | TRBJ1-5 |  | 52 | 23 | 4.3E-03 |
| Singleton | TRAV41 | X | TRBV14 | TRBJ1-1 |  | 27 | 8 | 4.5E-03 |
| Singleton | TRAV14DV4 | TRAJ54 | TRBV18 | X |  | 25 | 7 | 4.7E-03 |
| Singleton | TRAV13-1 | TRAJ30 | TRBV7-3 | X |  | 17 | 4 | 6.1E-03 |
| Singleton | TRAV36DV7 | TRAJ23 | X | TRBJ1-4 |  | 10 | 1 | 6.2E-03 |
| Singleton | TRAV8-3 | TRAJ20 | X | TRBJ2-5 |  | 77 | 41 | 6.5E-03 |
| Singleton | TRAV23DV6 | TRAJ41 | TRBV5-1 | X |  | 21 | 6 | 6.7E-03 |
| Singleton | TRAV1-1 | X | TRBV30 | TRBJ2-7 |  | 48 | 21 | 7.7E-03 |
| Singleton | X | TRAJ27 | TRBV2 | TRBJ2-4 |  | 19 | 5 | 8.3E-03 |
| Singleton | X | TRAJ40 | TRBV10-3 | TRBJ1-5 |  | 27 | 9 | 8.6E-03 |
| Singleton | X | TRAJ45 | TRBV12-5 | TRBJ2-1 |  | 25 | 8 | 8.7E-03 |
| Singleton | TRAV14DV4 | X | TRBV20-1 | TRBJ1-5 |  | 121 | 73 | 8.7E-03 |
| Singleton | TRAV25 | X | TRBV27 | TRBJ1-2 |  | 50 | 22 | 9.1E-03 |
| Singleton | TRAV24 | X | TRBV19 | TRBJ2-7 |  | 50 | 22 | 9.6E-03 |
| Singleton | X | TRAJ53 | TRBV24-1 | TRBJ1-5 |  | 24 | 7 | 1.1E-02 |
| Singleton | TRAV6 | TRAJ22 | TRBV14 | X |  | 13 | 2 | 1.1E-02 |
| Singleton | TRAV9-2 | TRAJ53 | TRBV9 | X |  | 55 | 25 | 1.1E-02 |
| Singleton | TRAV4 | TRAJ9 | TRBV13 | X |  | 11 | 2 | 1.3E-02 |
| Singleton | TRAV14DV4 | TRAJ39 | TRBV24-1 | X |  | 15 | 3 | 1.4E-02 |
| Singleton | TRAV13-2 | X | TRBV2 | TRBJ1-5 |  | 64 | 32 | 1.5E-02 |
| Singleton | TRAV22 | TRAJ28 | TRBV5-8 | X |  | 12 | 2 | 1.7E-02 |
| Singleton | TRAV22 | TRAJ28 | TRBV5-4 | X |  | 12 | 2 | 1.7E-02 |
| Singleton | TRAV27 | TRAJ39 | X | TRBJ1-2 |  | 45 | 20 | 2.2E-02 |
| Singleton | X | TRAJ7 | TRBV15 | TRBJ2-7 |  | 21 | 6 | 2.4E-02 |
| Singleton | TRAV12-1 | TRAJ45 | TRBV18 | X |  | 26 | 8 | 2.7E-02 |
| Singleton | TRAV1-1 | TRAJ13 | X | TRBJ1-2 |  | 42 | 19 | 2.8E-02 |
| Singleton | TRAV25 | X | TRBV19 | TRBJ2-1 |  | 81 | 44 | 2.9E-02 |
| Singleton | TRAV29DV5 | X | TRBV16 | TRBJ1-1 |  | 11 | 2 | 3.1E-02 |
| Singleton | TRAV41 | TRAJ43 | TRBV18 | X |  | 19 | 5 | 3.1E-02 |
| Singleton | TRAV21 | TRAJ18 | TRBV7-3 | X |  | 11 | 2 | 3.4E-02 |
| Singleton | X | TRAJ45 | TRBV2 | TRBJ2-1 |  | 72 | 38 | 3.4E-02 |
| Singleton | TRAV8-2 | X | TRBV14 | TRBJ2-1 |  | 44 | 19 | 3.4E-02 |
| Singleton | TRAV8-4 | X | TRBV14 | TRBJ2-1 |  | 44 | 19 | 3.5E-02 |
| Singleton | TRAV12-2 | TRAJ13 | TRBV25-1 | X |  | 13 | 2 | 3.5E-02 |
| Singleton | TRAV2 | TRAJ13 | TRBV28 | X |  | 37 | 15 | 4.1E-02 |

## References

[1] T. Mora and A. M. Walczak, “Quantifying lymphocyte receptor diversity,” bioRxiv, p. 046870, Apr. 2016.

[2] J. Rossjohn, S. Gras, J. J. Miles, S. J. Turner, D. I. Godfrey, and J. McCluskey, “T Cell Antigen Receptor Recognition of Antigen-Presenting Molecules,” Annu. Rev. Immunol., vol. 33, no. 1, pp. 169–200, 2015.

[3] A. K. Sewell, “Why must T cells be cross-reactive?,” Nat. Rev. Immunol., vol. 12, no. 9, pp. 669–677, 2012.

[4] T. P. Arstila, A. Casrouge, V. Baron, J. Even, J. Kanellopoulos, and P. Kourilsky, “A direct estimate of the human alphabeta T cell receptor diversity,” Science, vol. 286, no. 5441, pp. 958–961, Oct. 1999.

[5] B. Howie et al., “High-throughput pairing of T cell receptor α and β sequences,” Sci. Transl. Med., vol. 7, no. 301, p. 301ra131, Aug. 2015.

[6] E. S. Lee, P. G. Thomas, J. E. Mold, and A. J. Yates, “Identifying T Cell Receptors from High-Throughput Sequencing: Dealing with Promiscuity in TCRα and TCRβ Pairing,” PLOS Comput. Biol., vol. 13, no. 1, p. e1005313, Jan. 2017.

[7] “Datasets,” 10x Genomics. [Online]. Available: https://www.10xgenomics.com/resources/datasets/. [Accessed: 08-Jun-2019].

[8] T. Dupic, Q. Marcou, A. M. Walczak, and T. Mora, “Genesis of the alpha beta T-cell receptor,” PLOS Comput. Biol., vol. 15, no. 3, p. e1006874, Mar. 2019.

[9] J. A. Carter et al., “T-cell receptor αβ chain pairing is associated with CD4+ and CD8+ lineage specification,” bioRxiv, p. 293852, Apr. 2018.

[10] H. M. Berman et al., “The Protein Data Bank,” Nucleic Acids Res., vol. 28, no. 1, pp. 235–242, Jan. 2000.

[11] M. Scutari, “Bayesian Network Constraint-Based Structure Learning Algorithms: Parallel and Optimized Implementations in the bnlearn R Package,” J. Stat. Softw., vol. 77, no. 1, pp. 1–20, Mar. 2017.

[12] M.-P. Lefranc et al., “IMGT unique numbering for immunoglobulin and T cell receptor variable domains and Ig superfamily V-like domains,” Dev. Comp. Immunol., vol. 27, no. 1, pp. 55–77, Jan. 2003.

[13] J. Foote and A. Raman, “A relation between the principal axes of inertia and ligand binding,” Proc. Natl. Acad. Sci., vol. 97, no. 3, pp. 978–983, Feb. 2000.

[14] M. Shugay et al., “VDJdb: a curated database of T-cell receptor sequences with known antigen specificity,” Nucleic Acids Res., vol. 46, no. D1, pp. D419–D427, Jan. 2018.

[15] L. C. Garner, P. Klenerman, and N. M. Provine, “Insights Into Mucosal-Associated Invariant T Cell Biology From Studies of Invariant Natural Killer T Cells,” Front. Immunol., vol. 9, 2018.

[16] M. Salio, J. D. Silk, E. Yvonne Jones, and V. Cerundolo, “Biology of CD1- and MR1-Restricted T Cells,” Annu. Rev. Immunol., vol. 32, no. 1, pp. 323–366, 2014.

[17] R. Reantragoon et al., “Antigen-loaded MR1 tetramers define T cell receptor heterogeneity in mucosal-associated invariant T cells,” J. Exp. Med., vol. 210, no. 11, pp. 2305–2320, Oct. 2013.

[18] H. Huang et al., “Select sequencing of clonally expanded CD8+ T cells reveals limits to clonal expansion,” Proc. Natl. Acad. Sci. U. S. A., vol. 116, no. 18, pp. 8995–9001, Apr. 2019.

[19] R. Gowthaman and B. G. Pierce, “TCRmodel: high resolution modeling of T cell receptors from sequence,” Nucleic Acids Res., vol. 46, no. W1, pp. W396–W401, Jul. 2018.

[20] M. V. Pogorelyy et al., “Exploring the pre-immune landscape of antigen-specific T cells,” Genome Med., vol. 10, no. 1, p. 68, 25 2018.

